# Astrocytes mobilize a broader repertoire of lysosomal repair mechanisms than neurons

**DOI:** 10.1101/2025.09.07.674666

**Authors:** Erin M. Smith, Natali L. Chanaday, Sandra Maday

## Abstract

Lysosomal damage impairs proteostasis and contributes to neurodegenerative diseases, yet cell-type-specific differences in lysosomal repair remain unclear. Using a neuron-astrocyte coculture system, we compared responses to lysosomal injury induced by a lysosomotropic methyl ester. Both neurons and astrocytes showed lysosomal damage, marked by galectin-3 recruitment to lumenal lysosomal β-galactosides, elevated lysosomal pH, and engagement of lysophagy receptors TAX1BP1 and p62. However, astrocytes showed a preferential recruitment of ESCRT repair machinery to damaged lysosomes. Additionally, the lysosomal membrane reformation pathway regulated by the RAB7-GAP, TBC1D15, was more robustly activated in astrocytes. By contrast, the PITT pathway, mediating lipid transfer between the ER and damaged lysosomes, was engaged in both cell types. Our data reveal a divergence in how neurons and astrocytes mobilize repair pathways to manage lysosomal damage. These data may reflect differences in lysosomal resilience between astrocytes and neurons and inform therapeutic strategies to correct lysosomal dysfunction in neurodegenerative diseases.

## INTRODUCTION

Lysosomes are key degradative organelles that maintain cellular proteostasis by breaking down proteins and organelles^1^. Neural cells, particularly long-lived neurons and astrocytes, are especially dependent on lysosomal integrity to support essential cellular functions and ensure survival. Notably, lysosomal dysfunction is a hallmark of several neurodegenerative diseases, including Alzheimer’s Disease (AD) and Parkinson’s disease (PD)^2,3^. Pathogenic proteins such as amyloid β, tau and α-synuclein (aSyn), can permeabilize lysosomal membranes which enables escape from lysosomes^4–9^. Once in the cytoplasm, these fibrils can convert endogenous copies of the protein to pathogenic conformations^4–9^. Pharmacologically inducing lysosomal membrane damage or impeding pathways responsible for membrane repair can exacerbate this process and accelerate the propagation of toxic protein species^4,7,9,10^. Despite these insights, how different cell types in the brain, such as neurons or astrocytes, respond to lysosomal membrane damage is poorly understood. Recent proteomic profiling has revealed that lysosomal protein composition varies across cell types, including those in the brain^11–14^, suggesting that lysosomes may have specialized properties depending on the cell type. In fact, lysosomes in pancreatic cancer cells exhibit unique properties that confer enhanced protection to membrane damage and support rapid cell growth^15,16^. Understanding how lysosomes differ between neurons and astrocytes, particularly in their resilience to membrane damage, may uncover the molecular basis for selective susceptibility to neurodegenerative insults across cell types and guide the development of targeted therapeutic strategies to mitigate neurodegeneration.

Permeabilization of the lysosomal membrane triggers a cascade of cellular responses to manage the damage^3,17,18^. Early responses attempt to repair membrane damage, while late stage responses eliminate damaged lysosomes by autophagy and are balanced with transcriptional responses to generate new lysosomes^3,17,18^. Small perforations in the lysosomal membrane rapidly activate ESCRT-mediated repair to restore membrane integrity. ESCRT complexes (ESCRT-0, I, II and III) are well established for their roles in targeting ubiquitinated membrane proteins to lysosomes for degradation via the formation of multivesicular bodies^19–22^. In the brain, ESCRT proteins and the ESCRT-binding protein ALIX are highly expressed^23–25^ and have critical roles in neuronal development and synaptic function^23–28^. Notably, mutations in CHMP2B are associated with neurodegenerative diseases, including frontotemporal dementia (FTD) and Amyotrophic Lateral Sclerosis (ALS)^29–31^. Recent work also implicates the ESCRTs as a critical defender of lysosome integrity^4,9^. In this pathway, calcium leakage through perforations in the lysosomal membrane is detected by the calcium-binding protein ALG-2/PDCD6. ALG-2 then directly recruits ALIX/PDCD6IP and the ESCRT-I component, TSG101^32–36^. ALIX and TSG101 act in parallel pathways to initiate assembly of the ESCRT-III complex (CHMPs 1-7 and IST1); ALIX directly recruits ESCRT-III components and TSG101 acts via the ESCRT-II component, EAP30^33–37^. The ESCRT-III complex then forms a filament spiral that collaborates with the AAA ATPase VPS4 to seal the damaged membrane, either by direct membrane scission or via membrane stabilization to allow for spontaneous resealing^19,22,38–41^. Blocking ESCRT recruitment by knockdown of ALIX or TSG101, or by chelating calcium, prolongs lysosomal damage and leads to cell death, highlighting the critical role of ESCRTs in lysosome recovery^33,34,36,37^.

Another strategy to restore the lysosomal membrane is via the shuttling of lipids between damaged lysosomes and the endoplasmic reticulum (ER), a process termed phosphoinositide-initiated membrane tethering and lipid transport (PITT). In this pathway, calcium efflux from damaged lysosomes promotes phosphatidylinositol-4 kinase type 2α (PI4K2A) accumulation onto damaged lysosomes, leading to local production of phosphatidylinositol-4-phosphate (PtdIns4P) ^42,43^. PtdIns4P then recruits oxysterol-binding proteins (OSBPs) and OSBP-related proteins (ORPs), proteins which also bind to integral membrane proteins on the ER (e.g. VAPA/B), thereby tethering damaged lysosomes to the ER to facilitate a counter transport of lipids^42,43^. OSBPs transport cholesterol from the ER to the lysosomal membrane in exchange for PtdIns4P, and ORPs transport phosphatidylserine (PS) from the ER to the lysosomal membrane in exchange for PtdIns4P^42,43^. Cholesterol may stabilize the lysosomal membrane and PS may recruit downstream effectors involved in repair^42–44^.

Lysosomal damage can also stimulate membrane reformation mediated by a RAB7 GTPase-activating protein (GAP), TBC1D15^45^. At later stages of lysosome damage, TBC1D15 is recruited to damaged lysosomes via coordinated binding with the lysosomal membrane protein LIMP2 and ATG8 proteins^45^. TBC1D15 may promote the assembly of machinery involved in autophagic lysosomal reformation (e.g., dynamin-2, clathrin, and kinesin) allowing for lysosomal tubulation and the generation of protolysosomes^45^. Combined, several mechanisms can be employed to respond to lysosomal membrane damage. But how neurons and astrocytes differentially engage these pathways remains poorly understood.

In this study, we use a primary mouse neuron-astrocyte coculture system to investigate how neurons versus astrocytes respond to lysosomal damage induced by L-leucyl-L-leucine methyl ester (LLOMe)^46,47^, with a focus on three key pathways: ESCRT-mediated membrane repair, TBC1D15-mediated membrane reformation, and the PITT pathway involved in lipid shuttling. We show functional evidence that LLOMe effectively induces lysosomal damage in both neurons and astrocytes based on accumulation of galectin-3, dissipation of lysosomal pH, and recruitment of lysophagy receptors TAX1BP1 and p62. Despite lysosomal damage occurring in both cell types, astrocytes but not neurons preferentially recruit components of the ESCRT-mediated repair and TBC1D15-mediated reformation pathways. By contrast, both neurons and astrocytes activate the PITT pathway in response to lysosomal damage. These data suggest that neurons and astrocytes activate a distinct complement of lysosomal repair pathways to manage lysosomal injury. These differences may render cell-type-specific vulnerabilities to pathophysiological stimuli that induce lysosomal damage.

## MATERIALS AND METHODS

### Reagents

Transgenic mice expressing GFP-LC3 (eGFP fused to the C-terminus of rat LC3B) were obtained from the RIKEN BioResource Research Center (RBRC00806; strain name B6.Cg-Tg(CAG-eGFP/LC3)53Nmi/NmiRbrc; GFP-LC3#53) and maintained as heterozygotes. All animal protocols were approved by the Institutional Animal Care and Use Committee at the University of Pennsylvania. Primary antibodies for immunofluorescence include antibodies against GFP (Aves Lab Inc., GFP-1020), MAP2 (Millipore Sigma, ab5622), β3 tubulin (R&D Systems, MAB1195), EEAT1/GLAST (abcam, ab416), S100β (Sigma, S2532), LAMP1 (abcam, ab25245), TAX1BP1 (abcam, ab176572), p62 (abcam, ab109012), CHMP2B (abcam, ab33174), ALIX (Biolegend, 634502), TBC1D15 (abcam, ab121396), IST1 (Proteintech, 19842-1-AP), CHMP2A (Proteintech, 10477-1-AP), PI4K2A (A kind gift of Dr. Pietro De Camilli, Yale University, New Haven, CT)^48^, and ORP9 (Santa Cruz, sc-398961). Secondary antibodies for immunofluorescence include goat anti-chicken Alexa Fluor 488 (Jackson ImmunoResearch, 103-545-155), goat anti-rabbit Alexa Fluor 488 (Invitrogen, A11034), goat anti-mouse Alexa Fluor 488 (Invitrogen, A11029), goat anti-rabbit Alexa Fluor 594 (Invitrogen, A11037), goat anti-mouse Alexa Fluor 594 (Invitrogen, A11032), goat anti-rat Alexa Fluor 647 (Invitrogen, A21247), goat anti-rabbit Alexa Fluor 647 (Invitrogen, A21245), goat anti-mouse Alexa Fluor 647 (Invitrogen, A32728), donkey anti-chicken Alexa Fluor 488 (Jackson ImmunoResearch, 703-545-155), donkey anti-mouse Alexa Fluor 594 (Invitrogen, A21203), donkey anti-rat Alexa Fluor 647 (Invitrogen, A48272). Reagents include Hoechst 33342 (Molecular Probes, H3570), LysoTracker Deep Red (Invitrogen, L12492) and L-Leucyl-L-Leucine methyl ester (hydrochloride) (Cayman Chemicals, 16008). Lentivirus expression plasmids designed on VectorBuilder include CAMKIIa(short)>eGFP/3xGGGGS/hGALS3[NM_001357678.2] (VB240205-1287tut), GFAP(short)>eGFP/3xGGGGS/hGALS3[NM_001357678.2] (VB240205-1286nsc), CAMKIIa(short)>mCHMP2B[NM_026879.3]/3xGGGGS/eGFP (VB240924-1386kbq), and GFAP(short)>mCHMP2B[NM_026879.3]/3xGGGGS/eGFP (VB240924-1387atc). Lentivirus shRNA plasmids designed on VectorBuilder include eGFP-U6>mChmp2b[shRNA#1] (VB240411-1458erh), eGFP-U6>mChmp2b[shRNA#2] (VB240411-1459she), eGFP-U6>mPdcd61p[shRNA#1] (VB240411-1460gch), eGFP-U6>mPdcd61p[shRNA#2] (VB240411-1462dus), eGFP-U6>mTbc1d15[shRNA#1] (VB240411-1463njf), eGFP-U6>mTbc1d15[shRNA#2] (VB240411-1464ggt) and eGFP-U6>Nontargeting[shRNA#1] (VB10000-0001mty). All VectorBuilder constructs were validated by DNA sequencing using Plasmidsaurus. Lentivirus plasmid pMDLg/pRRE was a gift from Didier Trono (Addgene plasmid # 12251; http://n2t.net/addgene:12251; RRID:Addgene_12251)^49^. CMV-VSV-G was a gift from Bob Weinberg (Addgene plasmid # 8454; http://n2t.net/addgene:8454; RRID:Addgene_8454)^50^. pRSV-Rev was a gift from Didier Trono (Addgene plasmid # 12253; http://n2t.net/addgene:12253; RRID:Addgene_12253)^49^.

### Primary Neuron and Astrocyte Coculture

Primary mouse hippocampal or cortical astrocytes (either GFP-LC3 transgenic or non-transgenic) were isolated from neonatal pups of postnatal day 0-1 (as described in Kulkarni et al. 2020)^51^. GFP-LC3 transgenic brain tissue was distinguished from non-transgenic brain tissue at the start of the dissection based on the presence or absence of GFP fluorescence, respectively. The isolation of GFP-LC3 transgenic or non-transgenic astrocytes was performed separately. Cortical or hippocampal astrocytes were grown in glial media (DMEM [Gibco, 11965-084] supplemented with 10% heat inactivated fetal bovine serum [Hyclone, SH30071.03], 2 mM Glutamax [Gibco, 35050-061], 100 U/ml penicillin and 100 µg/ml streptomycin [Gibco, 15140-122]) on 10 cm tissue culture plates at 37°C in a 5% CO_2_ incubator until ∼70-80% confluent. Astrocytes received a full media change the day after initial isolation and every ∼3-4 days thereafter; dishes were tapped vigorously before changing media to dislodge any microglia. Non-transgenic hippocampal or cortical neurons were isolated from embryos on embryonic day 15 (as described in Kulkarni et al. 2020)^51^. Non-transgenic brain tissue was distinguished from GFP-LC3 transgenic brain tissue at the start of the dissection based on the absence or presence of GFP fluorescence, respectively. Neurons were plated at 1.15 million cells per 6 cm tissue culture dish filled with 12 x 12 mm acid-washed Deckgläser coverslips or 5 x 18 mm acid-washed Deckgläser coverslips coated with 0.1 mg/ml poly-L-lysine [Sigma, P2636] for cortical neurons and 0.5 mg/ml poly-L-Lysine [Sigma, P2636] for hippocampal neurons. Neurons were plated in attachment media (MEM [Gibco, 11095-072] supplemented with 10% heat inactivated horse serum [Gibco, 16050-122], 1 mM sodium pyruvate [Gibco, 11360-070], 33 mM glucose [Sigma, G8769], and 37.5 mM NaCl) for 1-3 hours at 37°C in a 5% CO_2_ incubator. Coverslips were then transferred to individual wells of 12-well (for 12 mm coverslips) or 6-well (for 18 mm coverslips) plates containing maintenance media (Neurobasal Medium [Gibco, 21103-049] supplemented with 2% B-27 [Gibco, 17504-044], 33 mM glucose, 37.5 mM NaCl, 2 mM GlutaMAX, 100 U/ml penicillin, and 100 μg/ml streptomycin), and grown for 4-5 days at 37°C in a 5% CO_2_ incubator. On neuron DIV 4-5, GFP-LC3 transgenic or non-transgenic hippocampal or cortical astrocytes were dissociated from 10 cm tissue culture dishes using 0.05% trypsin-EDTA [Gibco, 15400-054]. Glial media was added to inactivate the trypsin and astrocytes were resuspended in coculture media (Neurobasal Medium supplemented with 2% B-27, 1% G-5 [Gibco, 17503012], 0.25% (0.5 mM) GlutaMAX, 100 U/ml penicillin, and 100 μg/ml streptomycin). Maintenance media was removed from the neuron culture and astrocytes (in coculture media) were then plated onto the neuron culture at a ratio of 1 astrocyte to 7.5 neurons. For all cocultures, cortical neurons were cultured with cortical astrocytes, and hippocampal neurons were cultured with hippocampal astrocytes. On DIV 1-2 of coculture, 2 µM of AraC (anti-mitotic drug; Sigma, C6645), was added to inhibit astrocyte growth. Cocultures were grown for 7-8 DIV (neuron were a total of DIV 11-13) in coculture media at 37°C in a 5% CO_2_ incubator. For monoculture astrocyte experiments, non-transgenic astrocytes were dissociated from a 10 cm tissue culture dish as described above and plated at density of 100,000-200,000 cells onto FluoroDishes (World Precision Instruments, FD35-100) coated with 0.5 mg/ml Poly-L-Lysine. Astrocytes were grown in Glial Media at 37°C in a 5% CO_2_ incubator until they reached ∼70% confluency at 5-7 DIV for performing experiments.

### HEK293T Culture and Lentivirus Production

HEK293T cells (ATCC, CRL-3216) were maintained in HEK Media (DMEM supplemented with 10% heat inactivated fetal bovine serum, 2 mM Glutamax, 100 U/ml penicillin and 100 µg/ml streptomycin) and grown on 10 cm tissue culture dishes at 37°C in a 5% CO_2_ incubator. Once HEK293T cells reached 70-80% confluency, they were dissociated from the dish using 0.05% trypsin-EDTA and plated onto new 10 cm tissue culture dishes; cells were passaged for a maximum of 16 passages. For lentivirus production, HEK293T cells were plated in 6-well plates at densities of 150,000-300,000 cells per well and grown at 37°C in a 5% CO_2_ incubator. HEK293T cells were grown for 1-2 days to reach a confluency of 30-40%. Once HEK293T cells were 30-40% confluent, each well of a 6-well plate was transfected with 0.5 µg each of the following lentiviral packaging plasmids: pCMV-VSV-G (Addgene, #8454), pMDLG/pRRE (Addgene, #12251), pRSV-REV (Addgene, #12253), and 1.5 µg of plasmid expressing fluorescently-tagged proteins or 1 µg of plasmid to express shRNA. FuGENE6 (F6-1000) reagent was used for the transfection following the manufacturer’s protocol. One day post transfection, HEK media was replaced with coculture media. One day later, coculture media containing lentivirus was then collected from each well and centrifuged at 2000 RPMs for 15 minutes to get rid of cellular debris. The supernatant was collected, aliquoted, and stored at −80°C until further use. The recombinant lentiviruses used are all third-generation lentiviruses and are replication incompetent. Protocols for working with recombinant lentiviruses were approved by the Institutional Biosafety Committee (under the Environmental Health and Radiation Safety division) of the University of Pennsylvania.

### Lentiviral Transduction of Neuron-Astrocyte Coculture and Monocultured Astrocytes

On DIV 3-4 of coculture, lentiviruses to express eGFP-Galectin-3 or CHMP2B-eGFP were added to cocultured neurons and astrocytes at a 1:10 dilution and incubated for 4-5 days until DIV 7-8 of coculture (Fig. 1g-v, Fig. S1b-d, Fig. 3k-m). Whenever possible, the same lentivirus batch was used for all replicates of an experiment. Experiments using constructs with neuron or astrocyte specific promoters were performed on separate coverslips of cocultures. For experiments validating antibody specificity, a 1:10 dilution of lentiviruses to express shRNAs was added to monocultured astrocytes at 20-30% confluency. For knockdown samples, two different lentiviruses expressing different targeting shRNA sequences were pooled. ShRNAs were expressed for 3 days before cells were fixed at 60-70% confluency (Fig. S2).

**Figure 1.**
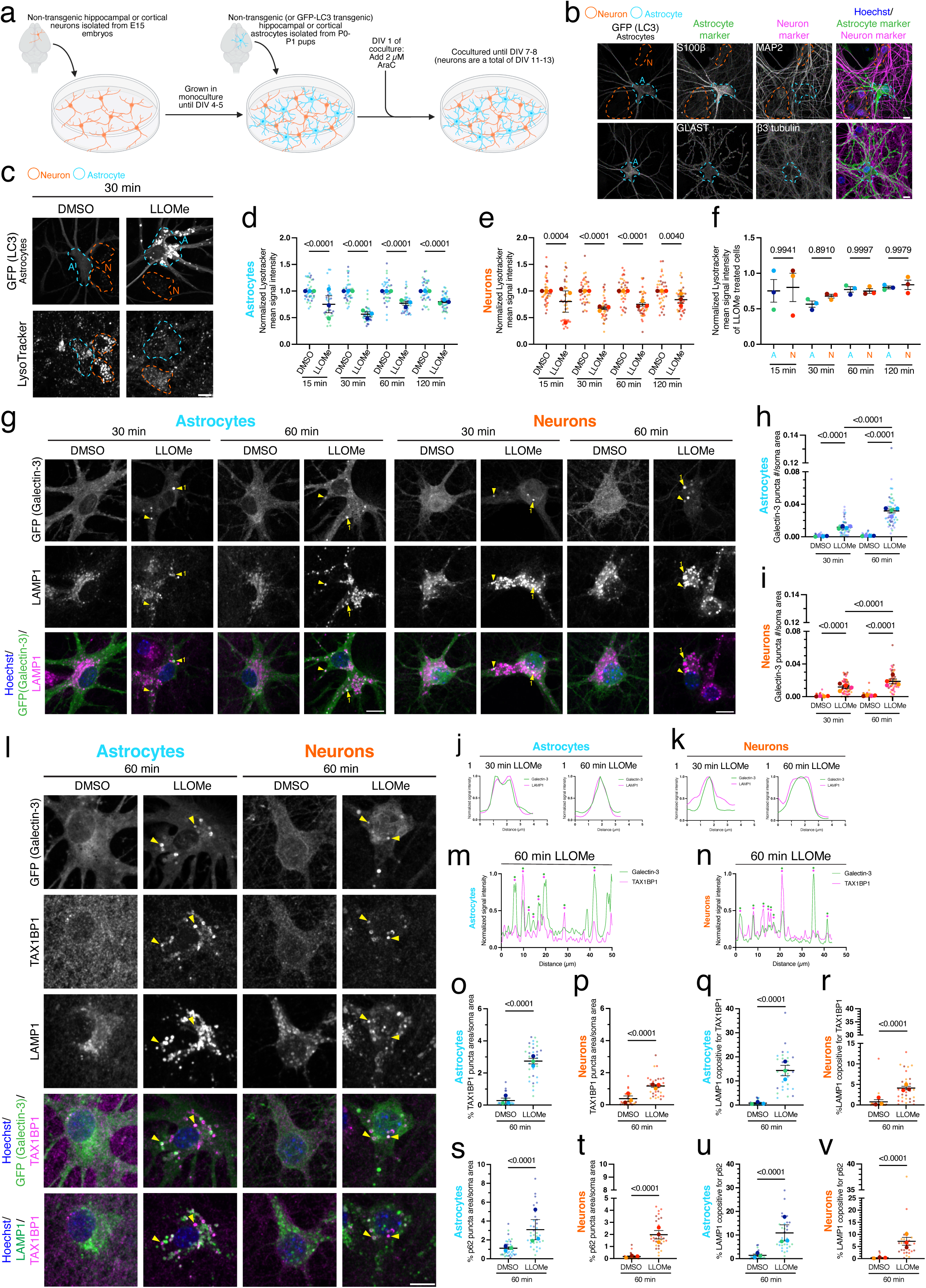
Lysosomotropic agent LLOMe effectively damages lysosomes in both astrocytes and neurons. **(a)** Schematic of neuron-astrocyte coculture. **(b)** Cocultured hippocampal astrocytes and neurons immunostained for GFP (LC3), astrocyte markers S100β or GLAST, and neuron markers MAP2 or β3 tubulin. **(c)** Cocultured hippocampal astrocytes and neurons treated with 1.5 mM LLOMe or equivalent volume of DMSO solvent control for 30 minutes and loaded with LysoTracker. **(d and e)** Corresponding quantification of normalized LysoTracker mean signal intensity with 1.5 mM LLOMe treatment as a function of time (15 to 120 minutes) in astrocytes **(d)** and neurons **(e)**. Horizontal bars represent the means of the biological replicates ± SEM. Shown are p-values from a Linear Mixed Effects (LME) model with Holm’s post hoc correction; N=35-38 astrocytes **(d)** and N=38-40 neurons **(e)** from 3 independent experiments. **(f)** Normalized mean signal intensity of 1.5 mM LLOMe treated astrocytes and neurons from **(d)** and **(e)** to enable direct comparison between cell types. Horizontal bars represent the means of the biological replicates ± SEM. Shown are p-values from a one-way ANOVA with Sidak’s multiple comparison correction; N=35-38 astrocytes and 38-40 neurons from 3 independent experiments. **(g)** Cocultured hippocampal astrocytes and neurons exogenously expressing eGFP-Galectin-3 were treated with 1.5 mM LLOMe or equivalent volume of DMSO solvent control for 30 or 60 minutes and immunostained for GFP (Galectin-3) and LAMP1. Yellow arrowheads indicate GFP (Galectin-3) puncta that are copositive for LAMP1. **(h and i)** Corresponding quantification of the number of GFP (Galectin-3) puncta normalized to cross-sectional soma area in LLOMe or DMSO treated astrocytes **(h)** and neurons **(i)**. Horizontal bars represent the means of the biological replicates ± SEM. Shown are p-values from an LME model with Holm’s post hoc correction; N=47-49 astrocytes **(h)** and N=51-55 **(i)** from 4 independent experiments. **(j and k)** Representative line scans of GFP (Galectin-3) and LAMP1 puncta from arrowheads labeled “1” in panel **(g)** from astrocytes **(j)** and neurons **(k)**. **(l)** Cocultured hippocampal astrocytes and neurons exogenously expressing eGFP-Galectin-3 were treated with 1.5 mM LLOMe or equivalent volume of DMSO solvent control for 60 minutes and immunostained for GFP (Galectin-3), TAX1BP1 and LAMP1. Yellow arrowheads indicate GFP (Galectin-3) puncta that are copositive for TAX1BP1 and LAMP1. **(m and n)** Representative lines scans of TAX1BP1 and GFP (Galectin-3) in LLOMe treated astrocytes **(m)** and neurons **(n)**; green and magenta dots indicate overlapping TAX1BP1 and Galectin-3 peaks. **(o and p)** Corresponding quantification of total TAX1BP1 puncta area normalized to soma area in LLOMe or DMSO treated astrocytes **(o)** and neurons **(p)**. Horizontal bars represent the means of the biological replicates ± SEM; shown are p-values from a LME model; N=33-36 astrocytes **(o)** and 32 neurons **(p)** from 3 independent experiments. **(q and r)** Corresponding quantification of total LAMP1 puncta area co-positive for TAX1BP1 in LLOMe or DMSO treated astrocytes **(q)** and neurons **(r)**. Horizontal bars represent the means of the biological replicates; shown are p-values from a LME model; N=30-35 astrocytes **(q)** and 30 neurons **(r)** from 3 independent experiments. **(s and t)** Corresponding quantification of data in Fig. S1b measuring total p62 puncta area normalized to soma area in astrocytes **(s)** and neurons **(t)**. Horizontal bars represent the means of the biological replicates ± SEM. Shown are p-values from a LME model; N=33-34 astrocytes **(s)** and 32-36 neurons **(t)** from 3 independent experiments. **(u and v)** Corresponding quantification of data in Fig. S1b measuring total LAMP1 puncta area co-positive for p62 in LLOMe or DMSO treated astrocytes **(u)** and neurons **(v)**. Horizontal bars represent the means of the biological replicates ± SEM. Shown are p-values from a LME model; N=29-32 astrocytes **(u)** and 29-33 neurons **(v)** from 3 independent experiments. Throughout the figure, neurons are outlined in orange and astrocytes are outlined in blue. Scale bars, 10 µm. For all graphs in all figures (except graphs of line scans), small circles indicate the measurements from individual cells (e.g., the technical replicates) from each of the independent experiments; large circles indicate the corresponding biological means from each of the independent experiments (e.g., the biological replicates); independent experiments are color-coded.

### LLOMe Treatments

For LLOMe treatments in Fig. 1-5 and Fig. S1, S3b, S5, S6, cocultured neurons and astrocytes were incubated in coculture media supplemented with 1.5 mM LLOMe or equivalent volume of solvent control (DMSO) for 15, 30, 60 or 120 minutes, as indicated directly in the figure and corresponding legend. For LLOMe treatment in Fig. S3a, cocultured neurons and astrocytes were incubated in coculture media supplemented with 750 µM LLOMe or equivalent volume of solvent control (DMSO) for 30 minutes. For antibody validation experiments in Fig. S2, monocultured astrocytes were incubated in glial media supplemented with 1.5 mM LLOMe for 30 minutes.

### Immunofluorescence

Following treatments, cells (either cocultured astrocytes and neurons or monocultured astrocytes) were fixed with 4% PFA/4% sucrose (PFA, Sigma, P6148; D-sucrose, Fisher Scientific, BP220-1) for 10 minutes, then washed 3 times with 1X PBS (150 mM NaCl, 50 mM NaPO_4_, pH 7.4) (as in Figs. 1-5 and Figs. S1-3, 5, 6). Cells were permeabilized with either 0.1% Triton X-100 (Fisher BioReagents, BP151-100) in 1X PBS for 5 minutes (as in Figs. 1-5, Figs. S1-3, 5), 0.1% Saponin (Sigma, S4521) in 1X PBS for 10 minutes (as in Fig. S6a-f), or 100% ice cold methanol (Fisher Scientific, A412-4) at −20°C for 10 minutes (as in Fig. S6g-l). Cells were then washed twice with 1X PBS and blocked for 1 hour at room temperature in goat block (1X PBS supplemented with 5% goat serum [Sigma, G9023], 1% BSA [Fisher BioReagents, BP1605], and 0.05% sodium azide [Fisher BioReagents, BP922I]) or horse block (1X PBS supplemented with 5% heat inactivated horse serum, 1% BSA, and 0.05% sodium azide). Horse block was used only when primary antibodies generated in mice or rats were used together for the immunostain (as in Fig. 5f-l). Goat block was used for all other experiments. Primary antibodies diluted in goat or horse block were then added to cells for 1 hour at room temperature. Cells were then washed 3 times for 5 minutes each with 1X PBS. Samples were then incubated in secondary antibodies diluted in goat or horse block for 1 hour at room temperature. Secondary antibodies raised in goats were used with the goat block and secondary antibodies raised in donkeys were used with the horse block. Samples were then washed 3 times for 5 minutes each with 1X PBS and rinsed one time with MilliQ water. Finally, coverslips were mounted in ProLong Gold (Thermo Fisher Scientific/Molecular Probes, P36930) and cured overnight at room temperature. Monocultured astrocytes on FluoroDishes were left in the third 1X PBS wash and imaged the same day. All steps of the immunostain procedure were performed with the samples protected from ambient light.

### Microscope

Both immunofluorescence and live-cell imaging for all experiments was performed on a BioVision spinning disk confocal microscope system with a Leica DMi8 inverted widefield microscope, a Yokagawa W1 spinning disk confocal, and a Photometrics Prime 95B scientific complementary metal–oxide–semiconductor camera. The microscope is equipped with an environmental chamber to maintain 37°C for live-cell imaging. Images were acquired with the VisiView software using a 63 ×/1.4 NA plan apochromat oil-immersion objective or a 40x/1.3 NA plan apochromat oil-immersion objective and solid state 405 nm, 488 nm, 561 nm and 647 nm lasers. Acquisition parameters were kept constant across treatment conditions and independent biological replicates.

### Cell Selection During Image Acquisition

During image acquisition, imaging frames were selected based on morphological criteria of astrocytes using an astrocyte-specific marker (e.g. GFP-LC3 transgene, or endogenous S100β or GLAST) and not the main markers of interest. Astrocytes selected for imaging had a stellate morphology with a minimum of three primary branches (often many more) and had a multitude of finer processes branching off of the primary branches. Astrocyte morphology shown in each figure is representative of the types of astrocytes that were selected for imaging and analysis. Given the higher ratio of neurons to astrocytes in the coculture, every frame selected based on the astrocyte also contained multiple neurons. For imaging of CHMP2B-eGFP and eGFP-Galectin-3, the marker used to assess cellular morphology was also the main marker of interest. However, neurons and astrocytes were still both selected for imaging based on morphological criteria. Astrocytes were selected using the criteria described above, and neurons were selected based on the presence of multiple well-developed dendrites and an axon.

### Immunofluorescence Imaging

Immunofluorescence imaging was performed on the spinning disk confocal described above. Z-stacks with a step size 0.2 µm that spanned the entire depth of both neuron and astrocyte somas were acquired. All coverslips were imaged within 1 week of staining to prevent potential decay in signal.

### Live-cell Imaging

For live-cell imaging of LysoTracker (Fig. 1c-f, Fig. S1a), samples were either preincubated with 100 nM of LysoTracker for 30 minutes before adding 1.5 mM of LLOMe for 15 minutes, coincubated with 100 nM of LysoTracker and 1.5 mM of LLOMe for 30 minutes, or treated first with 1.5 mM of LLOMe for 60 minutes or 120 minutes before adding 100 nM of LysoTracker for the final 30 minutes. This staggered treatment and LysoTracker addition allowed for all coverslips to receive a 30-minute incubation with LysoTracker and either a 15, 30, 60 or 120 minute treatment of 1.5 mM LLOMe. For live-cell imaging of CHMP2B-eGFP (Fig. 3k-m), samples were incubated with 1.5 mM LLOMe for 30 minutes. All treatments were performed in coculture media. At the completion of each treatment time, coverslips were washed once with HibE imaging solution (Hibernate E [Brain Bits, HE-Lf] supplemented with 2% B-27 and 2 mM GlutaMAX). Coverslips were then placed in a Chamlide magnetic imaging chamber (BioVision Technologies, CM-B18) and imaged in HibE imaging solution. Hoechst was added to the imaging solution at 1 µg/ml to stain nuclei. Live-cell imaging was performed on the spinning disk confocal described above. Z-stacks with a step size of 0.2 µm were acquired throughout the entire depth of both the neuron and the astrocyte somas. Each coverslip was imaged for a maximum of 30 minutes in an environmental chamber at 37°C.

### Image Analysis

#### Quantification of Puncta Area

To determine puncta area of a protein of interest (Fig. 1o, p, s, t; Fig. 2b, c, g, h; Fig. 3b, c, g, h; Fig. 5i, j; Fig. S1h; Fig. S5b, d; Fig. S6b, c, e, f, h, i, k, l) maximum intensity projections of Z-stacks were generated in FIJI. Neuron and astrocyte somas were outlined, and the cross-sectional area of each soma was measured in FIJI. Whenever possible, astrocytes were selected from the cell-type marker channel (e.g. GFP-LC3 transgene, or endogenous S100β or GLAST), and neurons were selected from the LAMP1 and Hoechst channels. For analysis of CHMP2B-eGFP (Fig. 3l, m), and CHMP2B and ALIX in neurons in Fig. S6c, f, i, l when LAMP1 immunostain was not present, we used a FIJI Macro that randomizes the images in the dataset and assigns a numerical title to each image to blind the identity of the images prior to outlining the soma. Outlined somas were imported into Ilastik, a machine learning based program, that was trained to identify and segment puncta based on the intensity, edge gradients and texture of the signal. Separate Ilastik programs were used to analyze neuron versus astrocyte somas to account for cell-type-specific differences in signal intensity and the ratio of cytosolic to punctate signal. Within a cell type, the same Ilastik program was used for all replicates of a given experiment to enable direct comparisons. Ilastik segmentations were imported into a FIJI macro and analyzed using the “analyze particles” function to measure the total segmented area occupied by puncta of interest. Total puncta area of the protein of interest was then normalized to the respective soma area and expressed as a percentage. Measurements from individual cells were treated as technical replicates and averaged to obtain the corresponding biological mean for each independent experiment.

**Figure 2.**
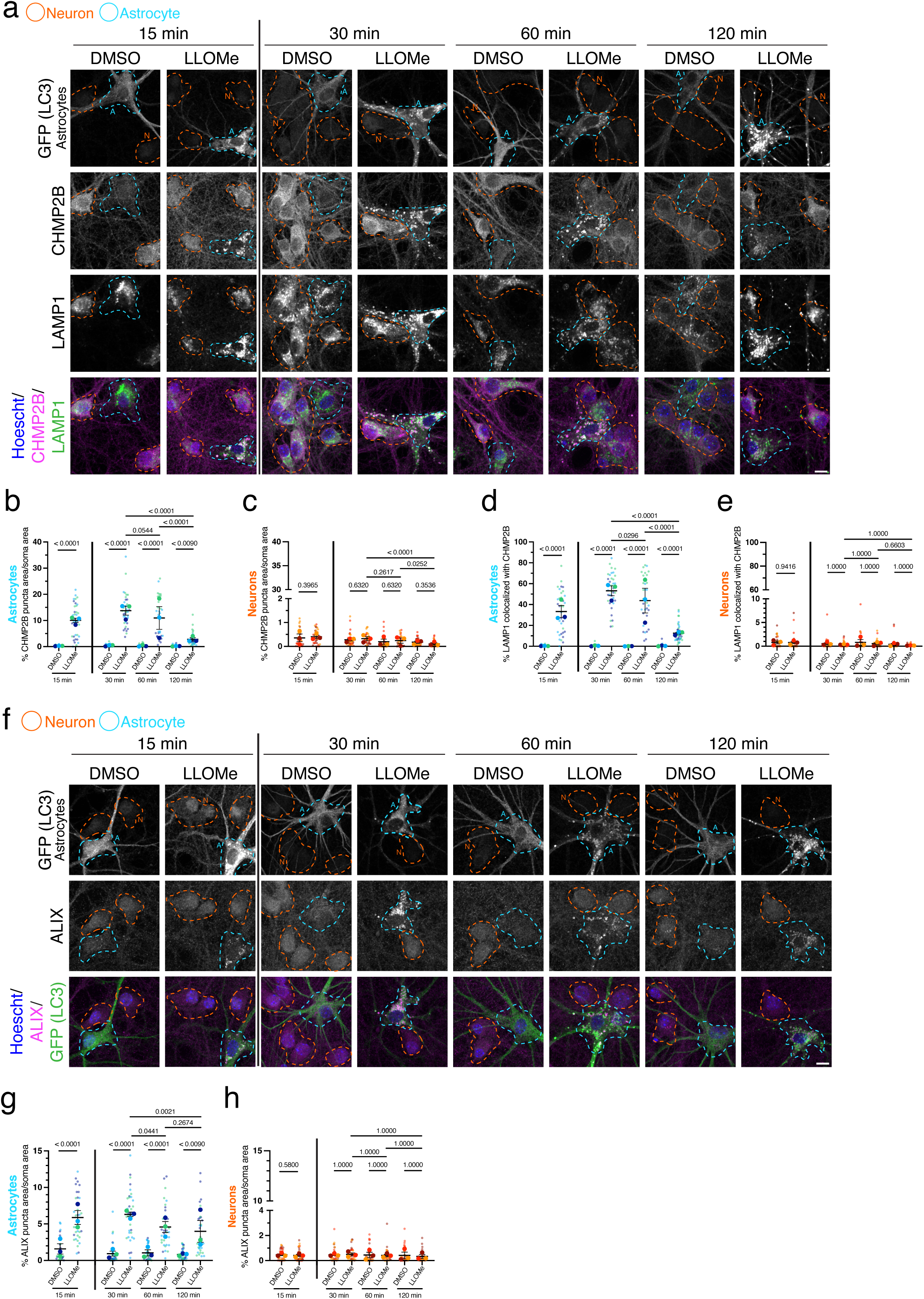
Damaged lysosomes preferentially recruit ESCRT components, CHMP2B and ALIX, in astrocytes as compared to neurons. **(a)** Cocultured hippocampal astrocytes and neurons treated with 1.5 mM LLOMe or equivalent volume of DMSO solvent control for the indicated time and immunostained for GFP (LC3), CHMP2B and LAMP1. **(b and c)** Corresponding quantification of total CHMP2B puncta area normalized to soma area in LLOMe or DMSO treated astrocytes **(b)** and neurons **(c)**. Line between 15 minutes and 30, 60 and 120 minutes denotes a separate set of experiments. For 15 minutes: horizontal bars represent the means of the biological replicates ± SEM; shown are p-values from a LME model; N=34-37 astrocytes **(b)** and N=34-36 neurons **(c)** from 3 independent experiments. For 30, 60 and 120 minutes: horizontal bars represent the means of the biological replicates ± SEM; shown are p-values from a LME model with Holm’s post hoc correction; N=28-37 astrocytes **(b)** and N=35-36 neurons **(c)** from 3 independent experiments. **(d and e)** Corresponding quantification of total LAMP1 puncta area co-positive for CHMP2B in LLOMe or DMSO treated astrocytes **(d)** and neurons **(e)**. Line between 15 minutes and 30, 60 and 120 minutes denotes a separate set of experiments. For 15 minutes: horizontal bars represent the means of the biological replicates ± SEM; shown are p-values from a LME model; N=33-35 astrocytes **(d)** and N=29-34 neurons **(e)** from 3 independent experiments. For 30, 60 and 120 minutes: horizontal bars represent the means of the biological replicates ± SEM; shown are p-values from a LME model with Holm’s post hoc correction; N=28-36 astrocytes **(d)** and N=30-36 neurons **(e)** from 3 independent experiments. **(f)** Cocultured hippocampal astrocytes and neurons treated with 1.5 mM LLOMe or equivalent volume of DMSO solvent control for the indicated time and immunostained for GFP (LC3) and ALIX. **(g and h)** Corresponding quantification of total ALIX puncta area normalized to soma area in LLOMe or DMSO treated astrocytes **(g)** and neurons **(h)**. Line between 15 minutes and 30, 60 and 120 minutes denotes a separate set of experiments. (For 15 minutes: horizontal bars represent the means of the biological replicates ± SEM; shown are p-values from a LME model; N=32-33 astrocytes **(g)** and N=38 neurons **(h)** from 3 independent experiments. For 30, 60 and 120 minutes: horizontal bars represent the means of the biological replicates ± SEM; shown are p-values from a LME model with Holm’s post hoc correction; N=34-37 astrocytes **(g)** and N=34-37 neurons **(h)** from 3 independent experiments. Throughout the figure, neurons are outlined in orange and astrocytes are outlined in blue. Scale bars, 10 µm.

### Colocalization Analysis for Proteins of Interest with LAMP1

To determine colocalization between two proteins (Fig. 1q, r, u, v; Fig. 2d, e; Fig. 3d, e, i, j; Fig. 5k, l; Fig. S1i), segmentation masks generated during puncta analysis were imported into a FIJI macro that uses the “and” function to compare the area of overlap between masks of two different channels from the same cell. To quantify the percentage of lysosomal area that is copositive for a protein of interest in each cell, the total area of overlap between LAMP1 and the protein of interest was normalized to the total puncta area for LAMP1 and plotted as percentage. Measurements from individual cells were treated as technical replicates and averaged to obtain the corresponding biological mean for each independent experiment.

**Figure 3.**
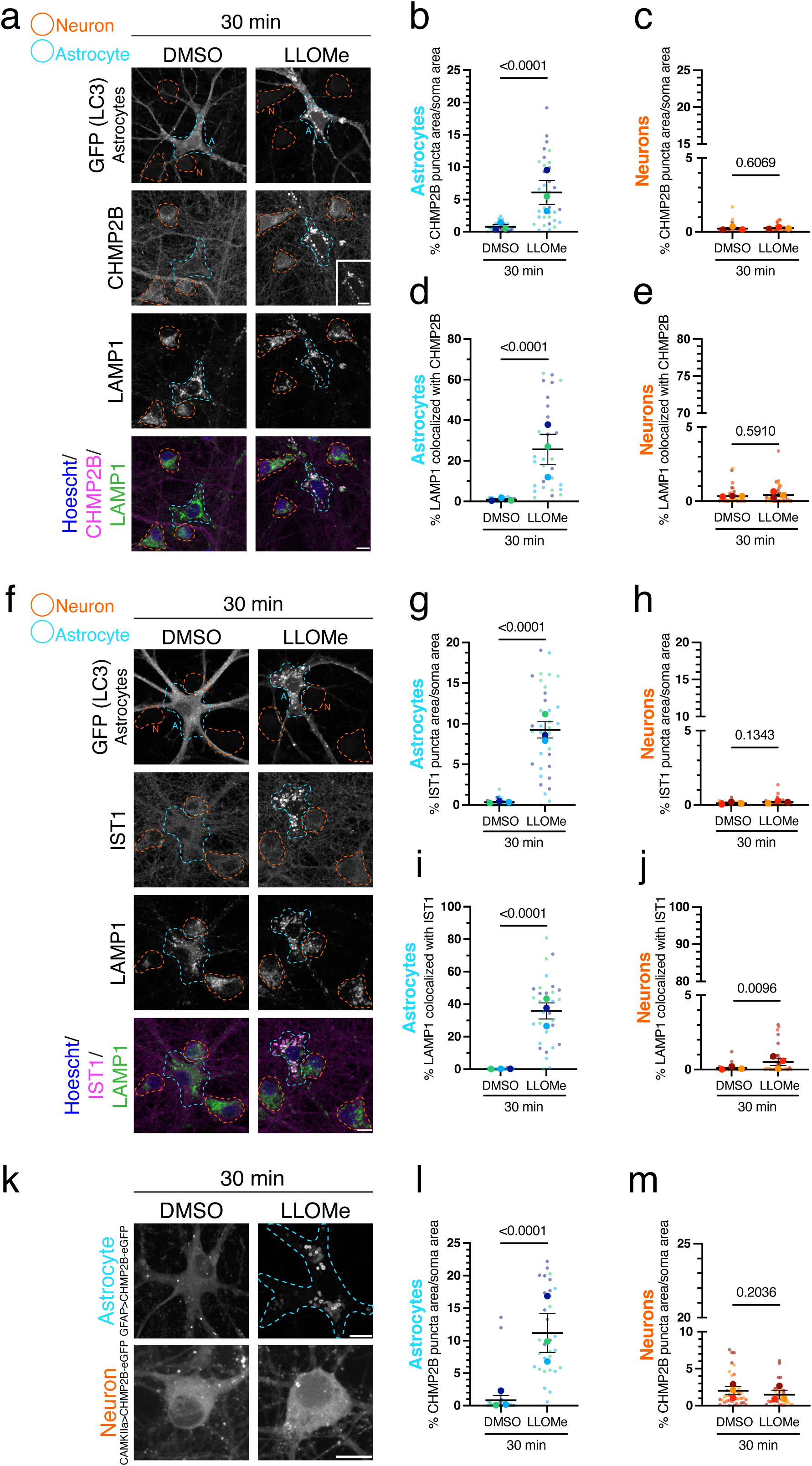
Preferential recruitment of ESCRT machinery to damaged lysosomes in astrocytes is conserved in astrocytes isolated from the mouse cortex. **(a)** Cocultured cortical astrocytes and neurons treated with 1.5 mM LLOMe or equivalent volume of DMSO solvent control for 30 minutes and immunostained for GFP (LC3), CHMP2B and LAMP1. Note: Inset showing astrocyte CHMP2B with LLOMe treatment is scaled to show CHMP2B-positive rings in astrocytes. **(b and c)** Corresponding quantification of total CHMP2B puncta area normalized to soma area in LLOMe or DMSO treated astrocytes **(b)** and neurons **(c)**. Horizontal bars represent the means of the biological replicates ± SEM. Shown are p-values from a LME model; N=29-34 astrocytes **(b)** and N=35-37 neurons **(c)** from 3 independent experiments. **(d and e)** Corresponding quantification of total LAMP1 puncta area copositive for CHMP2B in LLOMe or DMSO treated astrocytes **(d)** and neurons **(e)**. Horizontal bars represent the means of the biological replicates ± SEM. Shown are p-values from a LME model; N=27-30 astrocytes **(d)** and N=35 neurons **(e)** from 3 independent experiments. **(f)** Cocultured cortical astrocytes and neurons treated with 1.5 mM LLOMe or equivalent volume of DMSO solvent control and costained for GFP (LC3), IST1 and LAMP1. **(g and h)** Corresponding quantification of total IST1 puncta area normalized to soma area in LLOMe or DMSO treated astrocytes **(g)** and neurons **(h)**. Horizontal bars represent the means of the biological replicates ± SEM. Shown are p-values from a LME; N=29-35 astrocytes **(g)** and N=33-36 neurons **(h)** from 3 independent experiments. **(i and j)** Corresponding quantification of total LAMP1 puncta area copositive for IST1 in LLOMe or DMSO treated astrocytes **(i)** and neurons **(j)**. Horizontal bars represent the means of the biological replicates ± SEM. Shown are p-values from a LME model; N=25-33 astrocytes **(i)** and N=33-34 neurons **(j)** from 3 independent experiments. **(k)** Cocultured cortical astrocytes and neurons exogenously expressing CHMP2B-eGFP were treated with 1.5 mM LLOMe or equivalent volume of DMSO solvent control for 30 minutes. **(l and m)** Corresponding quantification of total CHMP2B-eGFP puncta area normalized to soma area in LLOMe or DMSO treated astrocytes **(l)** and neurons **(m)**. Horizontal bars represent the means of the biological replicates ± SEM. Shown are p-values from a LME model; N=31-39 astrocytes **(l)** and N=33-38 neurons **(m)** from 3 independent experiments. Throughout the figure, neurons are outlined in orange and astrocytes are outlined in blue. Scale bars, 10 µm.

### Quantification of LysoTracker Intensity

For LysoTracker imaging (Fig. 1c-f, Fig. S1a), maximum intensity projections of Z-stacks were generated in FIJI and neuron and astrocyte somas were outlined. For each soma, the mean intensity of the LysoTracker signal was measured in FIJI. For each independent experiment and cell type, LysoTracker mean intensities for DMSO at each timepoint were averaged. The LysoTracker mean intensities of both DMSO and LLOMe were then normalized to the corresponding DMSO average at each timepoint for each independent experiment. Normalized LysoTracker mean intensities for each soma (e.g. technical replicates) were averaged to generate a mean for each independent experiment (e.g. biological replicates) and plotted in PRISM. Statistical comparisons were only made within timepoints to account for normalization.

### Quantification of LAMP1 Puncta Integrated Density

To determine how LAMP1 puncta signal intensity changes with LLOMe treatments (Fig. S1f, g), maximum intensity projections of Z-stacks were generated in FIJI and neuron and astrocyte somas were outlined. LAMP1 puncta were identified and segmented as described above. Segmentations and soma outlines from each cell were imported into a FIJI macro that overlays the puncta segmentation onto the corresponding soma to measure the integrated density of all the puncta in the soma region. For each independent experiment and cell type, values for DMSO at each timepoint were averaged. The LAMP1 values for both DMSO and LLOMe were then normalized to the DMSO average at each timepoint for each independent experiment. Normalized LAMP1 integrated densities for each soma (e.g. technical replicates) were averaged to generate a mean for each independent experiment (e.g. biological replicates) and plotted in PRISM. Statistical comparisons were only made within timepoints to account for normalization.

### Quantification for Antibody Validation

For validating antibody specificity in monocultured astrocytes (Fig. S2), maximum intensity projections of Z-stacks were generated in FIJI and the entire astrocyte was outlined. The mean intensity of each marker of interest (e.g. CHMP2B, ALIX, and TBC1D15) was measured throughout the entire astrocyte using FIJI. For each independent experiment, the values for the non-targeting shRNA control were averaged. The values for the non-targeting shRNA control and the targeting shRNA were then normalized to the non-targeting shRNA average for each independent experiment. Normalized mean intensities for each astrocyte (e.g. technical replicates) were averaged to generate a mean for each independent experiment (e.g. biological replicates) and plotted in PRISM.

### Quantification of Galectin-3 Puncta Number

To determine Galectin-3 puncta number (Fig. 1h,i), maximum intensity projections of Z-stacks were generated in FIJI. Neuron and astrocyte somas were outlined, and the cross-sectional soma area was measured in FIJI. Soma outlines of the Galectin-3 stain were blinded using a FIJI macro that randomizes the images in the dataset and assigns a numerical title to each image. Using the cell-counter tool in FIJI, Galectin-3 puncta in each image were counted. Post-analysis, the identity of the data was decoded and the number of Galectin-3 puncta was normalized to the corresponding soma area (in µm^2^). Measurements from individual cells were treated as technical replicates and averaged to obtain the corresponding biological mean for each independent experiment.

### Colocalization Analysis Using Line Scans

To determine colocalization of Galectin-3 and LAMP1 (Fig. 1j, k), Galectin-3 and TAX1BP1 (Fig. 1m, n), Galectin-3 and p62 (Fig. S1c, d), GFP (LC3) and TBC1D15 (Fig. 4c), LAMP1 and TBC1D15 (Fig. 4e, f), PI4K2A and LAMP1 (Fig. 5b, c) and ORP9 and LAMP1 (Fig. 5g, h), a line scan analysis was performed in FIJI. A segmented line was drawn through puncta and cytosol in the Galectin-3 channel (Fig. 1j, k, m, n; S1c, d), the GFP(LC3) channel (Fig. 4c), the LAMP1 channel (Fig. 4e, f), the PI4K2A channel (Fig. 5b, c) and the ORP9 channel (Fig. 5g, h), and applied to the other marker of interest in the same location. Plot profile was used to measure intensity values of each pixel along the line for each marker. Intensity values were then normalized to the maximum intensity value within each respective marker. Normalized intensity values were plotted as a function of distance across the line and peaks that overlap were denoted with paired green and magenta dots.

**Figure 4.**
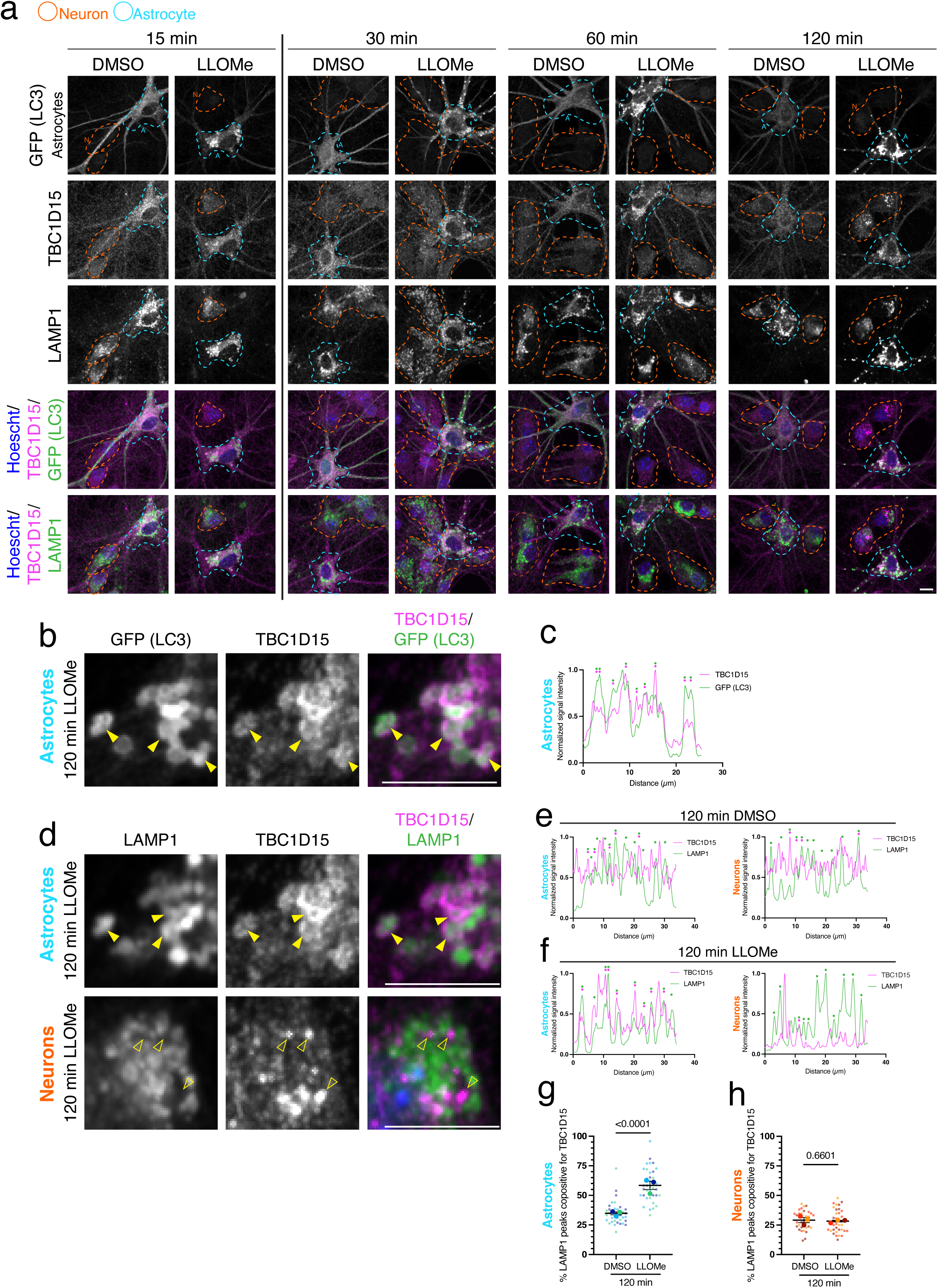
Rab7 GAP, TBC1D15, involved in lysosomal reformation is recruited to damaged astrocytic lysosomes. **(a)** Cocultured hippocampal astrocytes and neurons treated with 1.5 mM LLOMe or equivalent volume of DMSO solvent control for the time indicated and immunostained for GFP (LC3), TBC1D15 and LAMP1. **(b)** Zoom-in of an astrocyte treated with LLOMe for 120 minutes. Yellow arrowheads indicate GFP (LC3) and TBC1D15 overlap. **(c)** Representative line scan of TBC1D15 and GFP (LC3) in an astrocyte treated with LLOMe for 120 minutes. TBC1D15 and GFP (LC3) peaks that are overlapping are indicated with paired magenta and green dots. **(d)** Zoom-in of an astrocyte and neuron treated with LLOMe for 120 minutes. Closed yellow arrowheads indicate TBC1D15 puncta that overlap with LAMP1. Open yellow arrowheads indicate TBC1D15 puncta that do not overlap with LAMP1. **(e and f)** Representative line scans of TBC1D15 and LAMP1 puncta in neurons and astrocytes treated with DMSO for 120 minutes **(e)** or LLOMe for 120 minutes **(f)**. Paired magenta and green dots indicate overlapping TBC1D15 and LAMP1 peaks; green dots alone indicate LAMP1 peaks not overlapping with TBC1D15. **(g and h)** Corresponding quantification of total LAMP1 peaks copositive for TBC1D15 in LLOMe or DMSO treated astrocytes **(g)** and neurons **(h)**. Horizontal bars represent the means of the biological replicates ± SEM. Shown are p-values from a LME model; N=36-37 astrocytes (**g**) and N=33-34 neurons **(h)** from 3 independent experiments. Throughout the figure, neurons are outlined in orange and astrocytes are outlined in blue. Scale bars, 10 µm.

### Quantification of TBC1D15 and LAMP1 Line Scans

To further quantify the colocalization of TBC1D15 and LAMP1 (Fig. 4g, h), line scans were drawn in both cell type for each replicate. Normalized line scans were generated as described above. The number of total LAMP1 peaks and the number of LAMP1 peaks overlapping with TBC1D15 peaks were counted for each line scan. Only unambiguous peaks above background oscillations (e.g., above a normalized value of 0.25) were counted. The number of TBC1D15/LAMP1 overlapping peaks was divided by the total number of LAMP1 peaks and plotted as a percentage. Measurements from individual cells were treated as technical replicates and averaged to obtain the corresponding biological mean for each independent experiment.

### Figure Preparation and Statistical Analysis

All images shown in the figures are maximum projections generated from the Z-stack. For presentation of images, maximum and minimum gray values were adjusted linearly in FIJI. For a given experiment, the same adjustment was applied to all grayscale images within a marker across treatment conditions for a given cell type. However, in cases where lentiviruses were used to express exogenous proteins (e.g. eGFP-Galectin-3, CHMP2B-eGFP, or eGFP co-expressed with shRNAs), grayscale images were adjusted individually across treatment conditions due to variability in expression. Individual channels shown in merged images were also adjusted individually across treatment conditions to facilitate visualization of colocalization. Some panels showing astrocyte-specific markers were also adjusted individually across treatment conditions and denoted in the relevant supplementary figure legends. Graphs were plotted using GraphPad Prism. In all graphs in all figures (except graphs of line scans), small circles indicate the measurements from individual cells (e.g., the technical replicates) from each of the independent experiments; large circles indicate the corresponding biological means from each of the independent experiments (e.g., the biological replicates); bars are the means of the biological means ± SEM; independent experiments are color-coded. For Figure 1f, GraphPad Prism was used to perform a One-Way ANOVA with Sidak’s multiple comparison correction. For all other experiments, R studio version 2025.05.0.496 was used to perform either a linear mixed effects model (LME; R package “nlme”) or a LME model with the “weights” argument (R package “nlme”, weight = varIdent). In either case, a Holm’s post hoc correction was applied when multiple comparisons were made. For experiments in which time was not a factor (e.g., Fig. 1d, e, o-v; Fig. S1f, g; Fig. S2; Fig. 3; Fig 4; Fig. S5; Fig. S6) treatment condition was treated as a fixed effect, and biological replicate was treated as a random effect. For experiments in which time was a factor (e.g., Fig.1h, i; Fig. S1h, i; Fig. 2b-e, g, h; Fig. 5) the interaction of treatment and time was treated as the fixed effect, and biological replicate was treated as the random effect. Plots of residuals were used to assess homoscedasticity. For homoscedastic data, we used a LME model without weights (e.g., Fig. 1d, e, i, r, s, u; Fig. 2c, e, h; Fig. S2; Fig. 3c, e, h, l; Fig. 4; Fig. S5e-g; Fig. S6e, l). For non-homoscedastic data, we improved the LME model fit by using a LME model with the varIdent weights argument (e.g., Fig. 1h, o-q, t, v; Fig. S1; Fig. 2b, d, g; Fig. 3 b, d, g, i, j, m; Fig. 5; Fig. S5b, d; Fig. S6b, c, f, h, i, k). Images and graphs were assembled using Adobe Illustrator.

**Figure 5.**
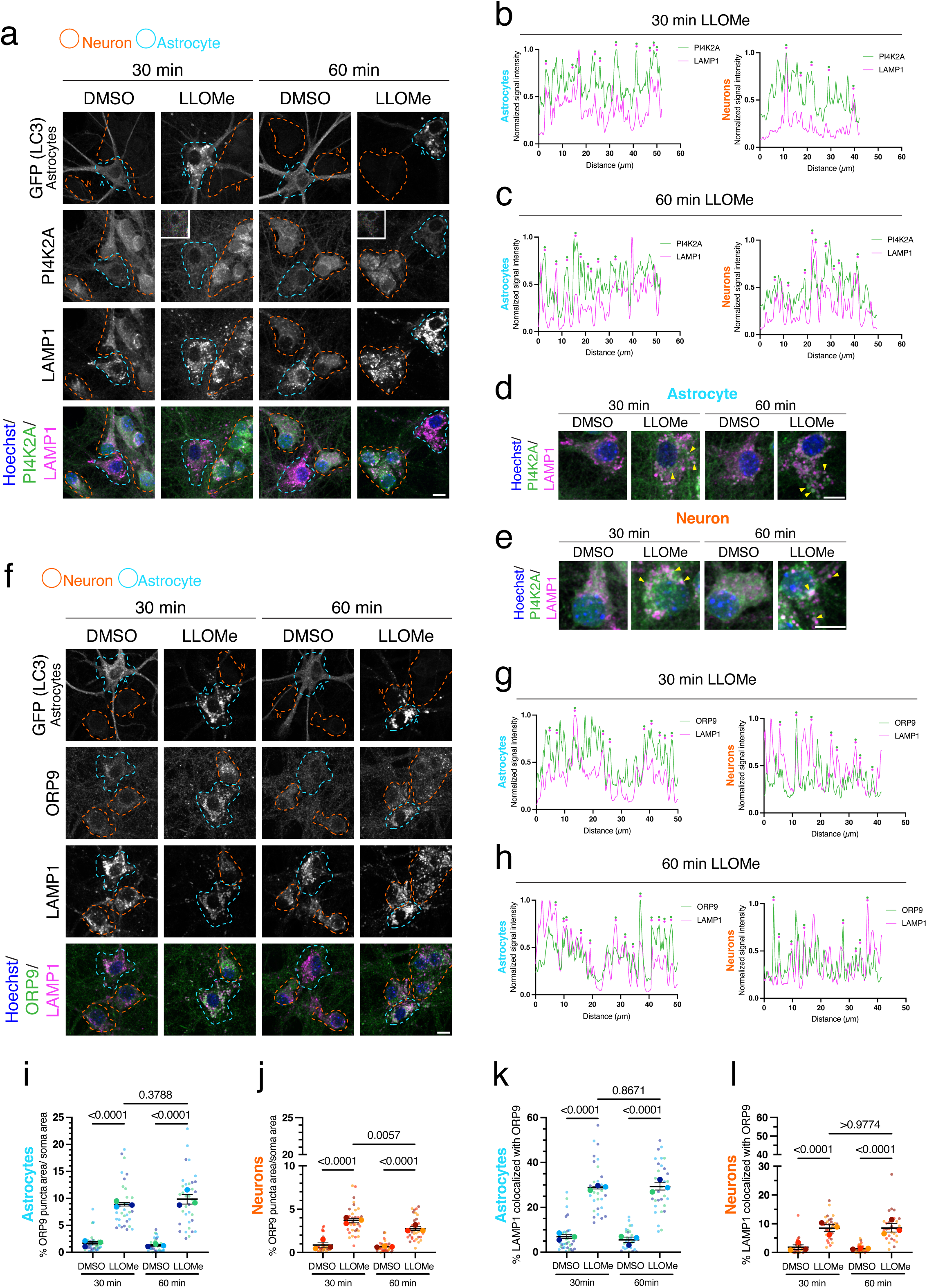
Machinery in the phosphoinositide signaling pathway that mediates lysosomal repair is recruited to damaged lysosomes in both astrocytes and neurons. **(a)** Cocultured hippocampal astrocytes and neurons treated with 1.5 mM LLOMe or equivalent volume of DMSO solvent control for the time indicated and immunostained for GFP (LC3), PI4K2A and LAMP1. Note: insets showing astrocyte PI4K2A with LLOMe treatment is enhanced to show PI4K2A rings in astrocytes. **(b and c)** Representative line scans of PI4K2A and LAMP1 puncta in astrocytes and neurons after 30 **(b)** and 60 **(c)** minutes of LLOMe. Overlapping peaks of PI4K2A and LAMP1 are denoted with paired green and magenta dots. **(d and e)** Zoom-ins of Hoechst/PI4K2A/LAMP1 merged images in LLOMe or DMSO treated astrocytes **(d)** and neurons **(e)**. Yellow arrowheads indicate PI4K2A puncta that are copositive for LAMP1 puncta. **(f)** Cocultured hippocampal astrocytes and neurons treated with 1.5 mM LLOMe or equivalent volume of DMSO solvent control for the time indicated and immunostained for GFP (LC3), ORP9 and LAMP. **(g and h)** Representative line scans of ORP9 and LAMP1 puncta in astrocytes and neurons after 30 **(g)** and 60 **(h)** minutes of LLOMe. Overlapping peaks of ORP9 and LAMP1 peaks are denoted by paired green and magenta dots. **(i and j)** Corresponding quantification of total ORP9 puncta area normalized to soma area in LLOMe or DMSO treated astrocytes **(i)** and neurons **(j)**. Horizontal bars represent the means of the biological replicates ± SEM. Shown are p-values from a LME model with Holm’s post hoc correction; N=33-36 astrocytes **(i)** and N=33-36 neurons **(j)** from 3 independent experiments. **(k and l)** Corresponding quantification of total LAMP1 puncta area copositive for ORP9 in LLOMe or DMSO treated astrocytes **(k)** and neurons **(l)**. Horizontal bars represent the means of the biological replicates ± SEM. Shown are p-values from a LME model with Holm’s post hoc correction; N=28-32 astrocytes **(k)** and N=28-31 neurons **(l)** from 3 independent experiments. Throughout the figure, neurons are outlined in orange and astrocytes are outlined in blue. Scale bars, 10 µm.

## RESULTS

### LLOMe effectively induces lysosomal damage in both neurons and astrocytes

To elucidate cell-type-specific responses of astrocytes and neurons to lysosomal damage, we cocultured primary mouse hippocampal or cortical astrocytes and neurons. In this system, neurons are cultured for 4-5 days *in vitro* to develop a dense meshwork of neurites, onto which astrocytes are plated and co-cultured with neurons for 7-8 days *in vitro* (neurons are cultured for a total of 11-13 days in vitro) (Fig. 1a). Direct contact with neurons induces a switch in astrocyte morphology from a polygonal form to more stellated and branched, adopting structures reminiscent of those observed *in vivo* (Fig. 1b). To visualize astrocytes in the coculture, we immunostained hippocampal cocultures for astrocyte-enriched markers S100β, a calcium binding protein, and GLAST, a glutamate transporter (Fig. 1b). S100β and GLAST reveal the elaborate and branched morphologies of astrocytes cocultured with neurons (Fig. 1b). Immunostain for neuron-specific cytoskeletal markers MAP2 to label dendrites and β3 tubulin to label neuronal microtubules reveal that neurons cocultured with astrocytes achieve highly polarized morphologies and a dense neurite meshwork (Fig. 1b). Thus, this system enables us to dissect molecular details of neuron-astrocyte biology.

To induce lysosomal damage, we treated cocultured hippocampal neurons and astrocytes with LLOMe, a lysosomotropic methyl ester that is processed by cathepsin C into membranolytic polymers that permeabilize the lysosomal membrane^46,47^. We validated that LLOMe effectively induced lysosomal damage in both neurons and astrocytes by quantifying a panel of lysosomal metrics. First, we measured the effect of LLOMe on a primary characteristic of active lysosomes, i.e. the maintenance of an acidic interior needed for optimal hydrolase function. We labeled acidic organelles using LysoTracker, a cell permeant dye that accumulates in acidic compartments. To distinguish astrocytes from neurons, hippocampal astrocytes were isolated from GFP-LC3 transgenic mice^52^ and co-cultured with non-transgenic hippocampal neurons (Fig. 1a). LC3 is a cytosolic protein that associates with autophagosome membranes^52^ but also provides a space fill to distinguish astrocyte morphology, as in Fig. 1b. We treated hippocampal neuron-astrocyte cocultures with 1.5 mM LLOMe for 15 min, 30 min, 60 min, and 120 min. Across all time points measured, LLOMe significantly reduced LysoTracker signal intensity in both neurons and astrocytes relative to the DMSO solvent control (Fig. 1c-e and S1a). To compare neurons and astrocytes directly, we quantified the average fold effect of LLOMe treatment for each time point across individual experiments (Fig. 1f). We found that the degree of reduction in LysoTracker signal with LLOMe treatment was not statistically different between neurons and astrocytes (Fig. 1f). We note that LysoTracker signal in the DMSO control was consistently higher in neurons versus astrocytes, but the fold reduction in LysoTracker signal with LLOMe was comparable between neurons and astrocytes (Fig. 1c and S1a). Thus, LLOMe reduces the presence of acidic organelles in neurons and astrocytes, consistent with increased lysosomal damage that neutralizes lysosomal pH. Moreover, LLOMe elicits a comparable effect on disrupting lysosomal acidity in both neurons and astrocytes.

Next, we investigated Galectin-3, a cytoplasmic lectin that binds to β-galactosides in the lumen of lysosomes that become exposed only during membrane damage^53,54^. Galectin-3 gains access to the interior of lysosomes with more extensive membrane damage and can facilitate their removal by autophagy, a process termed lysophagy^33^. Thus, Galectin-3 puncta are a measure of lysosomes with large membrane disruptions. To distinguish astrocytes and neurons, eGFP-Galectin-3 was expressed from a lentiviral vector using cell-type-specific promoters in cocultures of non-transgenic hippocampal neurons and astrocytes. To label neurons, we expressed eGFP-Galectin-3 using a neuron-specific CAMKIIa promoter and immunostained for GFP. In separate experiments, we labeled astrocytes by expressing eGFP-Galectin-3 using an astrocyte-specific GFAP promoter and immunostaining for GFP. Treatment with LLOMe (30 min or 60 min) significantly increased the number of Galectin3-positive puncta in the soma of both neurons and astrocytes relative to the DMSO solvent control (Fig. 1g-i). Importantly, Galectin-3 puncta colocalized with LAMP1, a marker for late endosomes and lysosomes (Fig. 1g, j, k). Only a subset of LAMP1 organelles were positive for Galectin-3 (Fig. 1g), highlighting the population of lysosomes with extensive membrane damage. We observed that LLOMe treatment increased the intensity and size of LAMP1-positive organelles resulting in the formation of large ring-like structures in both cell types (Fig. 1g), providing additional evidence of lysosomal damage. We also found that LLOMe-induced Galectin-3 puncta were present largely in the soma of neurons and astrocytes and less frequent in peripheral processes, consistent with the enrichment of mature and degradative lysosomes in the soma. In total, these data further support that LLOMe effectively causes ruptures in the lysosomal membrane that enables accessibility of Galectin-3 to lumenal lysosomal glycoproteins, in both neurons and astrocytes.

Lastly, we measured the recruitment of selective autophagy receptors to lysosomes in response to LLOMe in cocultured astrocytes and neurons. Sustained treatment with LLOMe initiates the selective elimination of damaged lysosomes by autophagy, termed lysophagy^3^. Lysophagy involves the recruitment of receptors (e.g., TAX1BP1 and p62/SQSTM1) that link ubiquitinated lysosomal membrane proteins to the autophagy machinery^3,55,56^. Thus, we immunostained hippocampal cocultures for TAX1BP1 or p62 and LAMP1; astrocytes or neurons were distinguished by the presence of eGFP-Galectin-3 expressed under a GFAP or CAMKIIa promoter, respectively. Treatment with LLOMe (60 min) elicited a striking increase in TAX1BP1 and p62 puncta in both neurons and astrocytes (Fig. 1I and S1b). Often, TAX1BP1 and p62 exhibited ring-like structures that localized to organelles co-positive for LAMP1 and Galectin-3 (Fig. 1I, m, n, S1b-d), indicating effective recruitment of lysophagy receptors to damaged lysosomes.

To quantify the recruitment of lysophagy receptors, we measured the total area of puncta positive for TAX1BP1 or p62 in neurons and astrocytes using Ilastik, a machine learning-based program to identify and segment objects of interest^57^. LLOMe treatment elicited a robust increase in total puncta area positive for TAX1BP1 or p62 in both neurons and astrocytes (Fig. 1o, p, s, t). Treatment with LLOMe also dramatically increased the percentage of LAMP1-positive puncta that were co-positive for TAX1BP1 or p62, in astrocytes and neurons (Fig. 1q, r, u, v). Thus, LLOMe increases the percentage of lysosomal organelles that recruit lysophagy receptors TAX1BP1 or p62 in both cell types. Quantification also revealed that LLOMe increased the intensity of LAMP1 signal in neurons and astrocytes (Fig. S1f, g), confirming our observations in Fig. 1g. To further determine lysosomal damage, we measured levels of GFP-LC3, a ubiquitin-like molecule involved in lysosomal damage pathways including lysophagy and conjugation of ATG8 to single membranes, (i.e., CASM)^58,59^. LLOMe treatment in astrocytes dramatically increased LC3-positive puncta and LC3 recruitment to lysosomal organelles (Fig. S1e, h, i), further confirming effective lysosomal damage. Taken together, multiple lines of functional evidence support effective lysosomal damage with LLOMe treatment in both neurons and astrocytes. Moreover, the damage can reach levels sufficient in both neurons and astrocytes to activate terminal mechanisms that will route lysosomes to autophagosomes for clearance.

### Lysosomal damage preferentially recruits ESCRT machinery in astrocytes as compared with neurons

The previous experiments validate the presence of late-stage responses to lysosomal damage that can clear unrepairable lysosomes (e.g. Galectin-3 and lysophagy receptors). Next, we wanted to investigate early-stage responses that function to repair damaged lysosomes. To elucidate how neurons and astrocytes initially respond to lysosomal damage, we measured the recruitment of ESCRT-mediated repair machinery, an early mechanism to repair small ruptures (∼100 nm) in the lysosomal membrane^36^. We treated hippocampal neuron-astrocyte cocultures with 1.5 mM LLOMe for 15 min and immunostained for CHMP2B, a core component of the ESCRT-III complex which forms the filament spiral needed for lysosome repair^19^. We first validated that the antibody against CHMP2B is specific by knocking down CHMP2B expression in monocultured astrocytes (Fig. S2a, b). In the coculture, astrocytes were distinguished from neurons by the presence of the GFP-LC3 transgene. In control cocultures, CHMP2B exhibits a higher expression level in neurons as compared with astrocytes, and localizes to punctate structures in the soma, consistent with established roles in MVB formation (Fig. 2a)^19^. After 15 min of LLOMe treatment, we observed a robust increase in CHMP2B-positive puncta in astrocytes (Fig. 2a). Importantly, CHMP2B localized to lysosomal organelles, often forming canonical ring-like structures decorating LAMP1-positive structures (Fig. 2a). CHMP2B-positive structures were also largely co-positive for GFP-LC3 (Fig. 2a). Interestingly, CHMP2B structures were predominantly localized to the soma of the astrocytes and proximal regions of astrocyte branches, similar to our observations with lysophagy receptors (Fig. 1I and S1b). By contrast, we did not observe an increase in CHMP2B puncta or a recruitment of CHMP2B to lysosomal organelles in neurons with 15 min of LLOMe treatment relative to the DMSO control (Fig. 2a). To determine whether this result is a consequence of slower repair kinetics in neurons, we measured CHMP2B levels at longer time points of LLOMe treatment. At 30, 60, and 120 min of LLOMe treatment, we continued to observe a preferential recruitment of CHMP2B to lysosomal organelles in astrocytes as compared with neurons (Fig. 2a). Thus, even with sustained LLOMe treatment, astrocytes more robustly recruit CHMP2B to damaged lysosomes as compared with neurons. Additionally, even a *lower* dosage of LLOMe (e.g. 750 µM LLOMe for 30 minutes) elicited the same effect (Fig. S3a).

Next, we quantified this effect by measuring the area occupied by CHMP2B. Across all time points, levels of CHMP2B puncta were elevated in astrocytes with LLOMe treatment as compared with the DMSO control (Fig. 2b). Moreover, CHMP2B levels peaked at early time points (e.g. 15 and 30 minutes) and began to decay at 120 minutes (Fig. 2b). Similarly, the percentage of LAMP1-positive organelles that colocalized with CHMP2B significantly increased at early time points and started to decay at 120 minutes (Fig. 2d). This temporal profile is consistent with prior reports showing ESCRT-mediated repair as an early response to lysosomal damage^36^, compared with more terminal lysosomal damage that recruits lysophagy machinery. By contrast, we did not observe a significant increase in the area of puncta positive for CHMP2B or the percentage of lysosomal organelles positive for CHMP2B in neurons treated with LLOMe versus DMSO at any of the time points investigated (Fig. 2c, e). These cell-type-specific differences are not attributed simply to differences in protein expression as CHMP2B protein appears to be higher in neurons versus astrocytes in control samples (Fig. 2a). Moreover, we mined the RNA sequencing database of neurons and glia generated by Zhang et al.^60^ to find that *Chmp2b* mRNA expression is also enriched in neurons as compared with astrocytes (Fig. S4a). Thus, protein levels of CHMP2B are likely not a limiting factor in neurons. We also examined the possibility that neurons may use the other isoform of CHMP2, CHMP2A; *Chmp2a* mRNA expression is also higher in neurons versus astrocytes (Fig. S4b)^60^. However, we found that astrocytes also preferentially recruit CHMP2A to damaged lysosomes as compared with neurons (Fig. S3b). LLOMe treatment induced ring-liked structures of CHMP2A that localized to LAMP1 in astrocytes, which were absent in neurons, albeit the signal for CHMP2A in astrocytes appeared more muted as compared with CHMP2B (Fig. S3b). In total, these data reveal an enhanced recruitment of ESCRT-mediated repair machinery to damaged lysosomes in astrocytes as compared with neurons.

We next investigated another component of the ESCRT-mediated repair machinery, ALIX. ALIX is recruited to damaged lysosomes via the calcium-binding protein ALG-2, and serves as a bridge to directly recruit ESCRT-III components to the lysosomal membrane^32–36^. We treated hippocampal neuron-astrocyte cocultures with 1.5 mM LLOMe for 15 min, 30 min, 60 min, and 120 min and immunostained for ALIX; astrocytes were identified by the GFP-LC3 transgene. We validated that the antibody against ALIX is specific by knocking down ALIX expression in monocultured astrocytes (Fig. S2c, d). Mining the database from Zhang et al.^60^, we validated that *Pdcd6ip* (ALIX) mRNA is expressed in both neurons and astrocytes at comparable levels (Fig. S4c). Further, the mRNA that encodes ALG-2/PDCD6 is also expressed in both neurons and astrocytes at comparable levels (Fig. S4d)^60^. Similar to our results with CHMP2B, we observed a robust increase in ALIX puncta preferentially in astrocytes as compared with neurons in response to LLOMe treatment, across all time points (Fig. 2f). We detected occasional ALIX puncta in the DMSO control in both neurons and astrocytes, but only astrocytes exhibited a dramatic increase in ALIX puncta with LLOMe (Fig. 2f). Moreover, in astrocytes, ALIX-positive puncta overlapped with GFP-LC3-positive structures (Fig. 2f). Quantification of ALIX puncta area revealed a similar temporal trend as we observed with CHMP2B (Fig. 2g). The levels of ALIX puncta in astrocytes peaked at 15 and 30 min of LLOMe treatment, and began to decline at 120 minutes (Fig. 2g). By contrast, we did not detect a significant increase in ALIX puncta in neurons across all time points investigated (Fig. 2h). Thus, we find that the differential recruitment of ESCRT repair machinery to damaged lysosomes in astrocytes versus neurons is not limited to CHMP2B, but is also observed with another component of this pathway, ALIX.

This enhanced recruitment of ESCRT-mediated machinery to damaged lysosomes in astrocytes versus neurons was very striking and unexpected. Thus, we performed additional controls to further assess these cell-type-specific differences. Our use of GFP-LC3 transgenic astrocytes served to distinguish astrocytes in the coculture from neurons, and in particular astrocytes with well-developed morphologies. However, emerging reports have since implicated ATG8 family members in new roles related to lysosomal repair via CASM^58,61–63^. Thus, we aimed to determine whether expression of exogenous GFP-LC3 was driving the recruitment of lysosomal repair machinery in astrocytes. For this experiment, we cocultured non-transgenic hippocampal neurons with non-transgenic hippocampal astrocytes and immunostained for CHMP2B or ALIX. Astrocytes were distinguished from neurons by immunostain for astrocyte-enriched markers S100β or GLAST (Fig. 1b). Cocultures were treated with 1.5 mM LLOMe for 30 minutes. Consistent with our prior observations, we observed a robust increase in CHMP2B and ALIX puncta area preferentially in astrocytes in response to LLOMe treatment (Fig. S5a-d). Importantly, the fold-increase in both CHMP2B and ALIX in non-transgenic astrocytes was comparable to our prior observations in GFP-LC3 transgenic astrocytes (Fig. 2b, g). As an additional control, the cross-sectional soma area of cocultured astrocytes that was used to normalize CHMP2B and ALIX signal is not affected by the LLOMe treatment and is also not contributing to the effect (Fig. S5e-g). Thus, this preferential recruitment of ESCRT-mediated repair machinery in astrocytes also occurs in non-transgenic astrocytes.

To check that these cell-type-differences in detecting ESCRT recruitment were not due to our sample preparation (i.e., permeabilization with 0.1% Triton X-100 for 5 minutes), we compared permeabilization with saponin versus methanol. We fixed using our standard protocol with paraformaldehyde but then permeabilized with either 0.1% Saponin for 10 minutes at room temperature or 100% methanol for 10 minutes at −20°C. These experiments were performed in hippocampal cocultures of non-transgenic neurons and non-transgenic astrocytes; astrocytes were identified by immunostain for GLAST or S100β. Cocultures were treated for 30 minutes with 1.5 mM LLOMe. With saponin permeabilization, we observed a robust increase in CHMP2B ring-like structures and ALIX puncta in astrocytes as compared with neurons in response to LLOMe treatment (Fig. S6a-f). With methanol, we also observed a preferential increase in CHMP2B rings and ALIX puncta in astrocytes as compared with neurons in response to LLOMe treatment (Fig. S6g-l). Thus, permeabilizing the cells with saponin or methanol shows similar results to our original protocol using Triton X-100, as evidenced by multiple markers of the ESCRT repair machinery (i.e., CHMP2B and ALIX). Therefore, the cell-type-specific differences in the recruitment of ESCRT machinery are not attributed to a difference in permeabilization procedures.

We next evaluated whether these differences in the ESCRT recruitment between neurons and astrocytes are limited to hippocampal cells or whether they are conserved in cells isolated from other brain regions. For this experiment, we generated cocultures of neurons and astrocytes isolated from the mouse cortex; cortical astrocytes were distinguished by GFP-LC3. We treated the cortical cocultures with 1.5 mM LLOMe for 30 minutes and immunostained for CHMP2B and IST1, a regulator of ESCRT-III assembly. Similar to hippocampal cocultures, LLOMe preferentially increases CHMP2B and IST1 puncta in cortical astrocytes as compared with cortical neurons (Fig. 3a-c, f-h). In cortical astrocytes, but not in cortical neurons, CHMP2B and IST1 form ring-like structures that colocalize with LAMP1 (Fig. 3a, d, e, f, i, j). Thus, these cell type differences in how neurons and astrocytes respond to lysosomal damage are conserved across different brain regions.

Lastly, we measured the recruitment of CHMP2B to damaged lysosomes in astrocytes and neurons using an orthogonal approach of live-cell imaging. We expressed CHMP2B-eGFP in non-transgenic cortical cocultures using a lentiviral vector and cell-type-specific promoters (i.e., CAMKIIa for neurons and GFAP for astrocytes). Measuring recruitment of exogenous fluorescently-labeled ESCRT subunits has been a standard approach in the field^36^. Cocultures were treated with 1.5 mM LLOMe for 30 minutes. In astrocytes treated with the DMSO control, CHMP2B-eGFP localized to the cytosol and to some puncta at baseline (Fig. 3k). With LLOMe treatment, CHMP2B-eGFP dramatically formed ring-like structures in the perinuclear region of astrocytes, reminiscent of structures observed by immunostain of endogenous CHMP2B (Fig. 3k). Corresponding quantification showed a dramatic increase in CHMP2B-positive structures in astrocytes with LLOMe (Fig. 3l). In neurons treated with the DMSO control, CHMP2B-eGFP localized to the cytosol and varying amounts of puncta, consistent with the distribution of endogenous CHMP2B (Fig. 3k). However, with LLOMe treatment, we did not observe a dramatic increase in puncta area or a redistribution to ring-like structures in neurons (Fig. 3k, m). Thus, with live-cell imaging of exogenous CHMP2B, we also detect a preferential enrichment of CHMP2B puncta in response to lysosomal damage in astrocytes but not in neurons. In sum, our data reveal a striking difference in how neurons and astrocytes respond to lysosomal damage.

### Lysosomal damage preferentially recruits TBC1D15, a Rab7-GAP involved in lysosomal membrane reformation in astrocytes as compared with neurons

Since neurons do not effectively recruit ESCRT repair machinery to damaged lysosomes, it is possible that they activate a complementary pathway. So next we measured the recruitment of TBC1D15 to damaged lysosomes in neurons and astrocytes. TBC1D15 is a Rab7-GAP involved in forming new lysosomes via membrane recycling of damaged lysosomes^45^. We treated hippocampal neuron-astrocyte cocultures with 1.5 mM LLOMe for 15 min, 30 min, 60 min, and 120 min and immunostained for TBC1D15; astrocytes were identified by the GFP-LC3 transgene. We validated that the antibody against TBC1D15 is specific by knocking down TBC1D15 expression in monocultured astrocytes (Fig. S2e, f). We also confirmed that *Tbc1d15* mRNA is expressed in both neurons and astrocytes (Fig. S4e)^60^. After 15 minutes of LLOMe treatment, we did not observe dramatic changes in TBC1D15 signal in astrocytes or neurons (Fig. 4a). However, by 30 minutes we began to see a redistribution of TBC1D15 to LAMP1-positive organelles in only astrocytes (Fig. 4a). By 60 and 120 minutes, we observed a robust enrichment of TBC1D15 signal on LAMP1-positive organelles in astrocytes (Fig. 4a, d). In fact, TBC1D15 formed rings around LAMP1-positive puncta (Fig. 4d). TBC1D15 rings also exhibited extensive overlap with GFP-LC3-positive rings (Fig. 4b, c). These ring structures are consistent with TBC1D15 decoration of the lysosomal membrane to facilitate membrane reformation events. After 120 minutes of LLOMe treatment, we did observe TBC1D15 puncta forming in neurons, however, these puncta largely did not overlap with LAMP1, suggesting that TBC1D15 was not recruited to damaged lysosomes (Fig. 4a, d). To quantify these differences in TBC1D15 localization to lysosomes, we generated line scans across LAMP1-positive puncta and counted the percentage of LAMP1 peaks that were co-positive for TBC1D15 peaks per cell (Fig. 4e, f). LLOMe treatment increased the percentage of lysosomes co-positive for TBC1D15 in astrocytes (Fig. 4e-g), but had no effect on LAMP1/TBC1D15 colocalization in neurons (Fig. 4e, f, h). Thus, these data suggest that astrocytes more effectively activate TBC1D15-mediated reformation pathways in response to lysosomal damage as compared with neurons.

### Astrocytes and neurons both recruit PITT pathway machinery in response to lysosomal damage

Thus far our data indicate that astrocytes but not neurons efficiently recruit lysosomal repair and reformation machinery. Another strategy that cells can employ to restore lysosome membrane integrity is via the transfer of lipids. Therefore, we examined the contributions of lipid shuttling via the PITT pathway to repair lysosomal membrane damage in neurons versus astrocytes^42,43^. We immunostained hippocampal cocultures for PI4K2A, the kinase that generates PtdIns4P on damaged lysosomes. The Zhang et al. database^60^ confirms expression of *Pi4k2a* mRNA in both astrocytes and neurons (Fig. S4f). In the DMSO control, we noted a reticular PI4K2A signal in neurons that resembled the Golgi (Fig. 5a), which is consistent with prior literature showing an enrichment of PI4K2A in the Golgi^64,65^. With LLOMe, we observed an increase in PI4K2A signal in neurons that appeared to surround LAMP1-positive puncta (Fig. 5a-c,e). In astrocytes, LLOMe treatment resulted in the formation of clear ring-like structures for PI4K2A, albeit the signal intensity was more muted as compared to the response in neurons (Fig. 5a-d). These data raise the possibility that lysosomal damage activates the PITT pathway in both astrocytes and neurons.

To more deeply explore the PITT pathway, we immunostained hippocampal cocultures for ORP9, which promotes membrane contact sites between damaged lysosomes and the endoplasmic reticulum to exchange PtdIns4P for PS. The Zhang et al. database^60^ confirms expression of *Osbpl9* (ORP9) mRNA in both astrocytes and neurons (Fig. S4g). We treated cocultures with 1.5 mM LLOMe for 30 min or 60 min; astrocytes were identified by the GFP-LC3 transgene. At each time point, we observed a robust increase in ORP9 puncta in astrocytes (Fig. 5f). Notably, we also observed a robust increase in ORP9 puncta in neurons (Fig. 5f). In both cell types, ORP9 signal was largely restricted to the soma and line scan analysis shows extensive colocalization of ORP9 with LAMP1 (Fig. 5g, h). Quantification revealed a robust increase in ORP9 puncta area (Fig. 5i, j), and in the percentage of LAMP1 puncta positive for ORP9 (Fig. 5k, l), in both neurons and astrocytes in response to LLOMe treatment. Thus, lysosomal damage robustly recruits ORP9, a key component of the PITT pathway, in both astrocytes and neurons.

Taken together, we found that lysosomal damage elicits a differential response in astrocytes versus neurons (Fig. 6, model). Astrocytes engage several pathways to mitigate lysosomal stress: ESCRT-mediated repair, TBC1D15-mediated reformation, and the PITT pathway. By contrast, neurons appear to activate only the PITT pathway, but not ESCRT-mediated repair or TBC1D15-mediated reformation. These cell-type-specific responses may confer selective vulnerabilities to neurodegenerative diseases caused by defects in lysosomal function.

**Figure 6.**
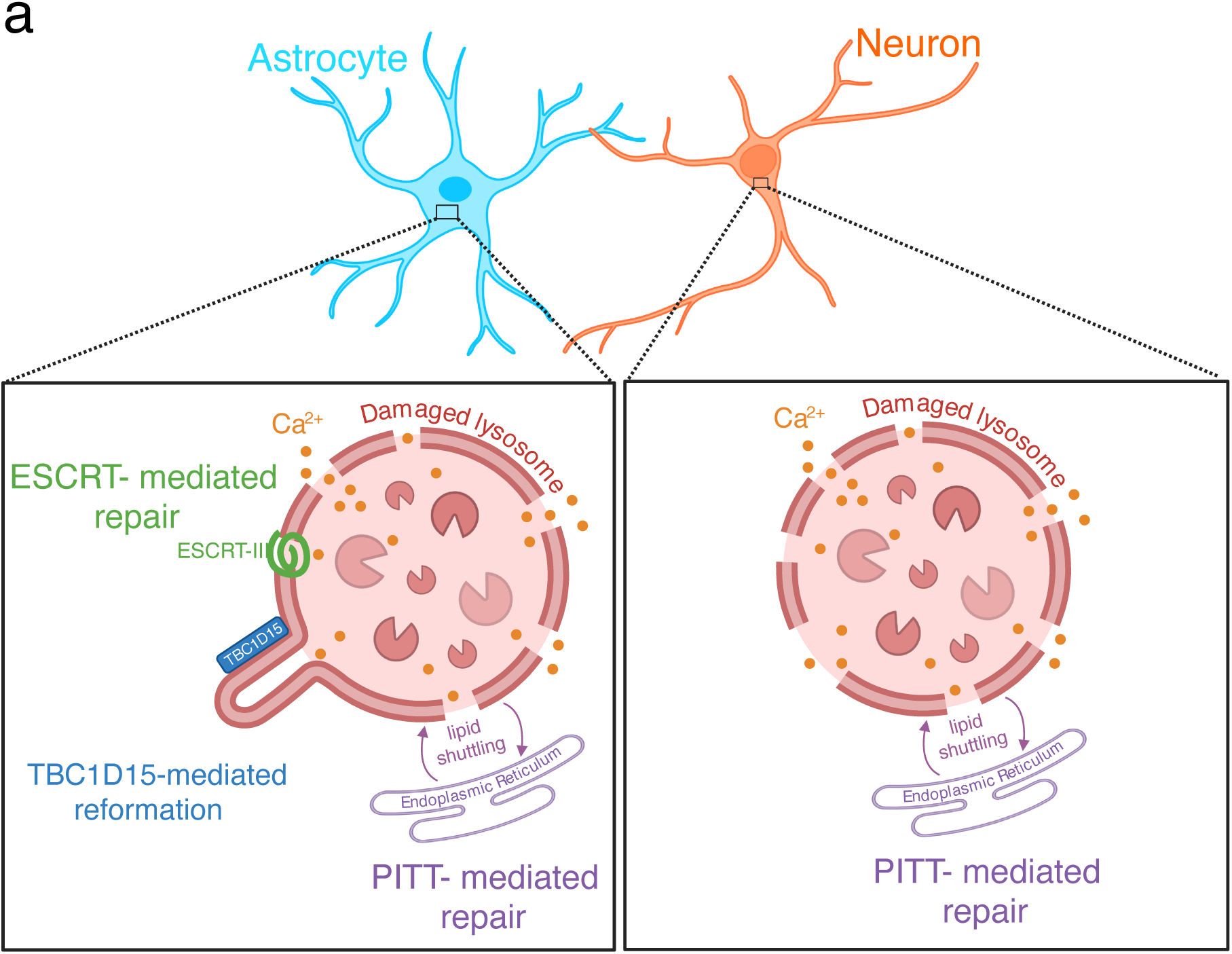
Model for the differential recruitment of lysosomal repair pathways in astrocytes versus neurons. In response to lysosome damage, astrocytes recruit the ESCRT pathway, the PITT pathway for lipid shuttling, and the TBC1D15-mediated reformation pathway. By contrast, neurons preferentially engage the PITT pathway.

## DISCUSSION

Here we report cell-type-specific responses to lysosomal damage in cocultured primary mouse hippocampal and cortical astrocytes and neurons. We find that sustained treatment with LLOMe recruits selective autophagy receptors p62 and TAX1BP1 to lysosomes in both cell types (Fig. 1I, S1b), activating terminal responses that eliminate damaged lysosomes via autophagy. However, shorter treatments with LLOMe elicit a differential early response in lysosomal repair in neurons versus astrocytes. Specifically, we analyzed the recruitment of three key lysosome repair/reformation pathways in neurons and astrocytes: (1) ESCRT-mediated repair, (2) TBC1D15-mediated pathway for lysosome membrane reformation, and (3) the PITT pathway to replenish lipids. We find that lysosomal damage recruits all three of these pathways in astrocytes, but only the PITT pathway in neurons (Fig. 2-5). We observed an enhanced recruitment of ESCRTs in astrocytes with not just one but multiple components of the pathway (e.g., CHMP2B, CHMP2A, ALIX, and IST1; Fig. 2, 3, S3, S5, S6). We also observed an enhanced recruitment of ESCRTs in astrocytes isolated from different brain regions (e.g., hippocampal and cortical astrocytes; Fig. 2, 3, S3). Lastly, we also consistently observed astrocyte-enhanced recruitment of ESCRTs across various experimental paradigms, regardless of GFP-LC3 expression (Fig. 3k-m, S5, S6), differing immunostain procedures (Fig. S6), and in both immunostaining of endogenous proteins or live-cell imaging of exogenous components (Fig. 3k-m). Mining the database by Zhang et al. shows that mRNA expression of key components of each repair pathway is comparable between both cell types (Fig. S4a-g)^60^. Notably, we find that CHMP2B protein expression under basal conditions appears higher in neurons as compared with astrocytes (Fig. 2a). These findings complement extensive literature showing key roles for ESCRT-III subunits and ALIX in proper neuronal development and synaptic function^23–27^, and evidence that CHMP2B variants are linked to the neurodegenerative disorders FTD and ALS^29,30^. Thus, the lack of robust ESCRT recruitment is not simply attributed to insufficient protein expression in neurons. Rather, neurons demonstrate a bias toward using the PITT pathway to repair damaged lysosomes. Thus, our data reveal a divergence in how neurons and astrocytes mobilize repair pathways to manage lysosomal damage.

These cell-type-specific responses in repair pathways may reflect differences in lysosomal resilience between astrocytes and neurons and confer a selective vulnerability to aging and neurodegenerative disease. Several studies implicate lysosomes as a point of vulnerability across multiple neurodegenerative diseases such as PD^66^ and AD^67–70^. Lysosomal rupture may be a conserved mechanism that enables amplification of pathogenic proteins within the brain^4–9^. In models of PD, fibrils or misfolded aSyn protein can rupture the lysosomal membrane and escape to the cytoplasm where they can corrupt the endogenous cytosolic protein to amplify fibril formation^6,71^. In models of AD, Tau fibrils can induce nanoscale perforations in the lysosomal membrane that can serve as nucleation points at the membrane surface for seeding soluble cytosolic protein^9^. Post-mortem brain tissue from AD patients shows elevated levels of lysosomal damage in the cerebral cortex that correlate with amyloid accumulation^72^. By contrast, Sanyal et al. propose that neuronal lysosomes may be naturally leaky which enables internalized fibrils of aSyn to gain access to the cytoplasm to poison the soluble aSyn^73^. Precisely how our observed differences in recruiting repair pathways affect lysosomal resilience and contribute to the amplification of pathogenic proteins remains an open question. The engagement of fewer repair pathways in neurons as compared with astrocytes may render neuronal lysosomes more susceptible to accumulating membrane perforations. Alternatively, the PITT pathway may be sufficient to effectively repair membrane damage, eliminating the need to activate other pathways. De Tito et al. have shown that PI4K2A is delivered from the Golgi to damaged lysosomes, via ATG9A vesicles^64^. We find that PI4K2A levels at baseline are higher in neurons as compared with astrocytes (Fig. 5a). Thus, perhaps neurons are more primed to shuttle PI4K2A to damaged lysosomes for rapid generation of PtdIns4P. Lastly, it also remains possible that astrocytic lysosomes may naturally experience more membrane damage and thus engage multiple pathways to restore lysosomal homeostasis. Future work will need to distinguish these possibilities.

Our study focuses on three key lysosome repair and reformation pathways, however future work will investigate cell-type-specific contributions of additional pathways in response to lysosomal membrane damage^17,18^. For example, stress granules are recruited to damaged lysosomes^74^ and they may facilitate repair by plugging small perforations and stabilizing the membrane^75^. Bonet-Ponce et al. report that, in astrocytes, the leucine-rich repeat kinase 2 (LRRK2) is recruited to damaged lysosomes to generate tubules, which may help to reroute material to other lysosomes^76^. A recent study found that VPS13C, a bridge-like lipid transport protein, localizes to ER-lysosome contact sites following damage which may facilitate the exchange of lipids independent of the PITT pathway^77^. Niekamp et al., find that sphingomyelin redistributes from the inner to the outer leaflet of damaged lysosomes, a mechanism important for repair that is triggered by calcium but independent of ESCRTs^78^. Loss of lysosome acidity or calcium efflux via activation of TRPML1, a cation channel localized to late endosomes and lysosomes, is sufficient to assemble the V_1_ and V_0_ subunits of the V-ATPase on the lysosomal membrane to promote reacidification of the lysosome^79,80^. Lastly, neurons and astrocytes may exhibit different thresholds for activating compensatory pathways for lysosome biogenesis to replace damaged lysosomes with new lysosomes. Thus, several pathways operate to restore and preserve lysosomal function that may also contribute to cell-type-specific differences in lysosomal resilience.

How is damage sensed in neurons versus astrocytes that might account for the differential recruitment in repair pathways? Lysosomes contain a high concentration of calcium (∼0.5 mM), at a level that is ∼5000-fold higher than cytosolic calcium (∼100 nM)^81^. Calcium release from the lysosome, that might be liberated via membrane perforations, has a central role in recruiting lysosomal repair pathways. Release of lysosomal calcium recruits ESCRTs to damaged lysosomes via the calcium-binding protein ALG-2^32,35^. ALG-2 recruits ALIX which coordinates assembly of ESCRT-III polymers^35^. In fact, in vitro reconstitution experiments using giant unilamellar vesicles demonstrates that calcium is sufficient to recruit ALG-2 to membranes and orchestrate downstream assembly of ALIX and ESCRT-III^35^. The involvement of calcium aligns with the rapid assembly kinetics observed in cells (occurring within minutes) in response to lysosomal damage^36^. At later time points of lysosomal damage, ESCRT retention on the lysosome may be influenced by interactions with GABARAPs^82^ and Galectin-3^33^, as described below. Calcium may also initiate the PITT pathway. In fact, activation of TRPML1, which would release calcium from the lysosome, is sufficient to localize PI4K2A to lysosomes^43^. The ability of calcium to activate both the ESCRT and PITT pathways suggests that presence of an additional regulatory layer that distinguishes cell type-specific responses.

Perhaps neurons have a higher degree of inter-organelle contacts that more effectively engages the PITT pathway. However, Rhoads et al. find that ER-Lysosome contacts might be slightly enriched in astrocytes as compared with neurons^83^. Interestingly, they find that neurons may exhibit more changes in inter-organelle contacts in response to cellular stress^83^. Chou et al. find that treatment of transdifferentiated neurons with LLOMe depolarizes mitochondria to the same degree as treating with the ionophore FCCP which uncouples oxidative phosphorylation in mitochondria^72^, raising the possibility that stress may propagate from lysosomes to mitochondria via contact sites. We observed an increase in TBC1D15 puncta in neurons with LLOMe, however, these puncta were largely distinct from lysosomes (Fig. 4a, d-h). Since TBC1D15 was also identified to localize to damaged mitochondria to coordinate engulfment by autophagosomes^84^, and neurons appear to have a higher density of mitochondria at baseline^83^, our data raise the possibility that TBC1D15 localized to mitochondria in neurons treated with LLOMe. Thus, stress may propagate differently in neurons versus astrocytes due to differences in organelle densities and contact sites, thereby activating different stress response pathways.

Galectins also play a key role in managing lysosomal damage. Galectin-3 localization to damaged lysosomes recruits ALIX and subsequent ESCRT-III effectors during acute damage, and coordinates autophagy induction during sustained damage^33^. Galectin-8 localization to damaged lysosomes leads to mTOR inactivation, facilitating a transcriptional response to generate new lysosomes^85^. Galectin-9 localization to damaged lysosomes facilitates ubiquitination of damaged lysosomes and activation of AMPK, a kinase that regulates autophagy levels^85,86^. Curiously, *Lgals3* (Galectin-3) mRNA expression is low in both astrocytes and neurons, but relatively lower in neurons^60^ (Fig. S4h). In corroboration, Galectin-3 protein expression in mouse primary cortical neurons or i^3^Neurons is undetectable relative to HeLa cells^56^. This relative expression pattern in neurons versus astrocytes might explain the enhanced recruitment of ESCRTs in astrocytes (Fig. 2-3, S5-6). *Lgals8* (Galectin-8) mRNA is expressed comparably in neurons and astrocytes^60^ (Fig. S4h). However, *Lgals9* (Galectin-9) mRNA is present at very low levels in both neurons and astrocytes^60^ (Fig. S4h), suggesting that some mechanisms identified in non-neuronal cells may not have a central role in the context of neurons and astrocytes. In models of neurodegeneration, Galectin-3 expression is increased in astrocytes and microglia, key regulators of inflammatory responses in the central nervous system^87^. Indeed, administration of Aβ peptides into the mouse hippocampus increases Galectin-3 in astrocytes and microglia, and increased Galectin-3 is coupled with cytokine production^88^. Galectin-3 expression is also upregulated in microglia in models of Huntington’s Disease and this elevation in Galectin-3 is causally linked to increased inflammation via the transcription factor NF-κB^89^. Interestingly, LLOMe activates NF-κB and subsequent expression of cytokines in cultured epithelial cells, suggesting that lysosome damage can trigger inflammatory pathways^90^. These data raise an intriguing possibility that lysosomal damage in astrocytes induces Galectin-3 expression which may amplify ESCRT mediated repair and also coordinate inflammatory responses.

Another signal that can recruit downstream effectors linked to lysosomal repair is CASM, a process that results in the lipidation of multiple ATG8 protein family members (e.g. LC3A, LC3B, LC3, GABARAP, GABARAPL1, GABARAPL2) directly onto lysosomal membranes^58,59^. CASM can (1) promote ESCRT association^82^, (2) facilitate recruitment of TBC1D15^45,84^, (3) recruit stress granule proteins which may help inactivate mTOR^74^, (4) recruit and activate LRRK2 on lysosomes^91^, and (5) stimulate TRPML1-mediated calcium efflux to activate TFEB^92^. Interestingly, most of these functions in lysosomal repair preferentially require GABARAP family members more than LC3 family members^45,74,82,84,91^. Thus, differences in lysosomal GABARAP conjugation may influence cell-type-specific differences in recruiting lysosomal repair pathways.

Improving lysosomal resilience may provide therapeutic value. In fact, stimulating calcium release from the lysosome via TRPML1 activation to preload the lysosome with ALG-2 and ESCRTs protects against membrane damage induced by osmotic stress^32^. Vest et al. identified a brain-penetrant small molecule C381 that targets the lysosome to promote lysosomal acidification and cargo degradation^93^. Treatment with C381 mitigates membrane permeabilization induced by LLOMe and also reduces lysosomal damage and proteotoxic burden observed in several neurodegenerative disease models^72,93^. Thus, pharmacological enhancement of lysosomal resilience can reverse proteostasis deficits and ameliorate disease pathology. Moreover, Pascua-Maestro et al. find that the neuroprotective effects of Apolipoprotein D (i.e., ApoD), an extracellular lipid-binding protein, include guarding the integrity of the lysosomal membrane against oxidative stress^94^. Upon oxidative stress, ApoD is endocytosed and concentrates on a subset of lysosomes where it reduces lipid peroxidation and membrane permeabilization^94^. Targeting other aspects of lysosome biology has also proven effective in mitigating disease pathology. In fact, Snead et al. identified a small molecule that can modulate the transport and distribution of lysosomes in neuronal axons, a process that is coupled with their maturation into degradative compartments, and resolve AD-linked pathologies in cultured human neurons^95^. Lastly, delivery of acidic nanoparticles to lysosomes can restore lysosomal acidification and degradation defects and prevent aSyn induced neurodegeneration in PD models^96,97^. Enhancing lysosomal degradative activity by delivery of recombinant pro-cathepsin D has also been proposed to reduce PD-linked pathologies^98^. Given the central role of lysosomal dysfunction in various neurodegenerative diseases, lysosomes represent a key therapeutic target. An exciting area of future investigation will be to assess the effects of enhancing lysosomal function and resilience in a cell-type-specific manner on disease pathology across several neurodegenerative models to help engineer more targeted therapeutic strategies.

## ACKNOWLEDGMENTS

This work was supported by NIH grants R01NS110716 and R21AG088697 to SM and a Synergy Grant from the Perelman School of Medicine at the University of Pennsylvania to SM. We thank Dr. Mary Putt (Biostatistics and Bioinformatics Core at the Intellectual and Developmental Disabilities Research Center at the Children’s Hospital of Philadelphia) and Dr. Edward Lee (Perelman School of Medicine at the University of Pennsylvania) for advice with statistical analyses. We are grateful to Dr. Phyllis Hanson (University of Michigan) for insightful discussions and helpful feedback on this study. We thank Dr. James Shorter (Perelman School of Medicine at the University of Pennsylvania) for constructive feedback on the manuscript.

## AUTHOR CONTRIBUTIONS

Conceptualization, E.M.S. and S.M.; Methodology, E.M.S., N.L.C, S.M.; Validation, E.M.S., S.M.; Investigation and Formal Analysis, E.M.S.; Resources, S.M.; Data Curation, E.M.S., S.M.; Writing – Original Draft, E.M.S., S.M.; Writing – Review and Editing, E.M.S., N.L.C, S.M.; Visualization, E.M.S., S.M.; Supervision, Project administration, and Funding Acquisition, S.M..

---

**Supplemental Figure 1.**
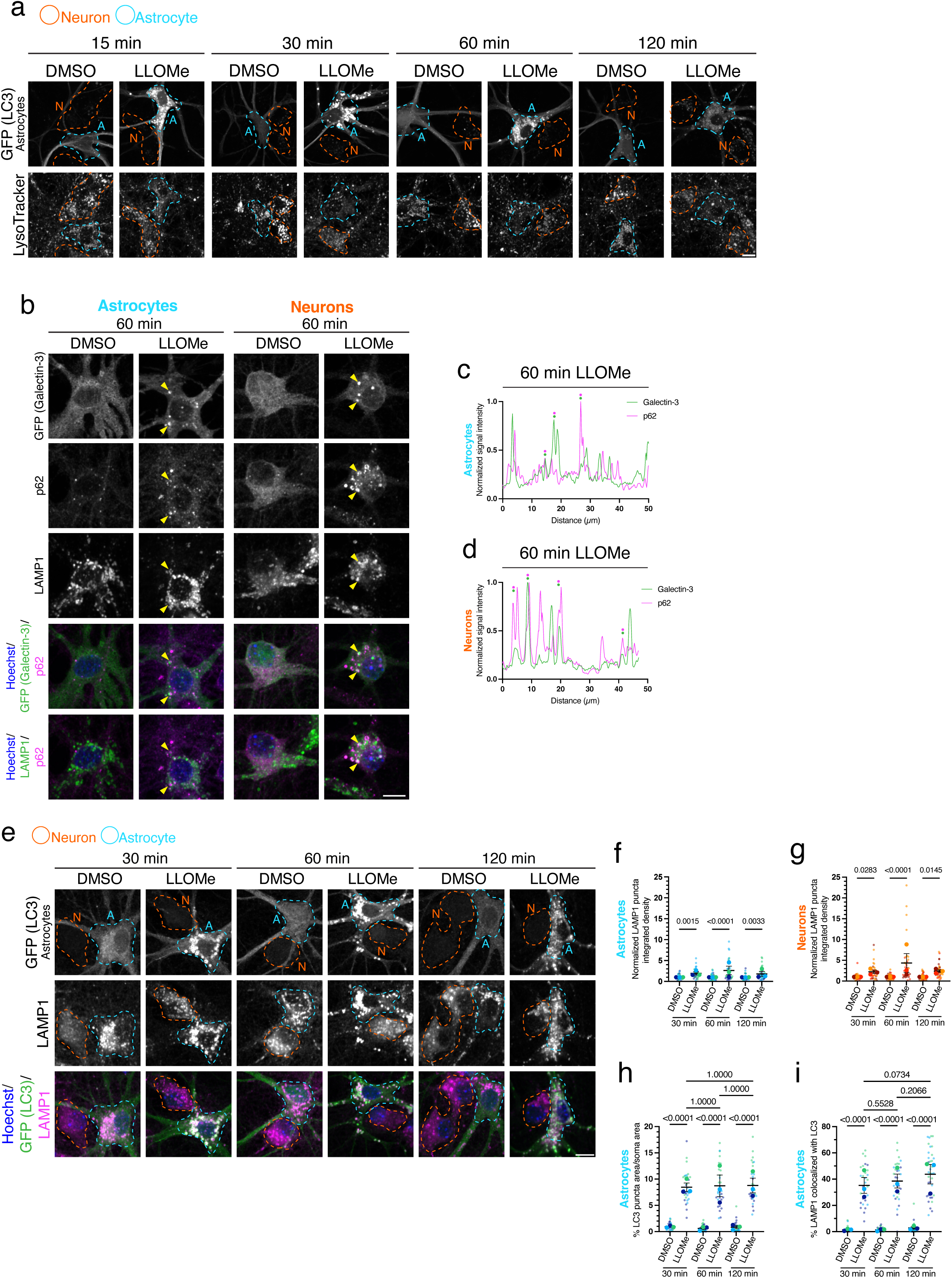
Additional metrics further support that LLOMe damages lysosomes in both astrocytes and neurons. **(a)** Cocultured hippocampal astrocytes and neurons treated with 1.5 mM LLOMe or equivalent volume of DMSO solvent control for the time indicated and loaded with LysoTracker. Images from the 30-minute timepoint are repeated from Figure 1c. **(b)** Cocultured hippocampal astrocytes and neurons exogenously expressing eGFP-Galectin-3 treated with 1.5 mM LLOMe or equivalent volume of DMSO solvent control and immunostained for GFP (Galectin-3), p62 and LAMP1. Yellow arrowheads indicate GFP (Galectin-3) puncta that are copositive for p62 and LAMP1. **(c and d)** Representative line scans of GFP (Galectin-3) and p62 puncta in astrocytes **(c)** and neurons **(d)** after 60 minutes of LLOMe treatment. Paired green and magenta dots indicate puncta positive for both markers. **(e)** Cocultured hippocampal astrocytes and neurons treated with 1.5 mM LLOMe or equivalent volume of DMSO solvent control for the indicated times and immunostained for GFP (LC3) and LAMP1. (**f and g)** Corresponding quantification of LAMP1 puncta normalized integrated density in astrocytes **(f)** and neurons **(g)** after LLOMe or DMSO treatment. Horizontal bars represent the means of the biological replicates ± SEM. Shown are p-values from a LME model with Holm’s post hoc correction; N=33-39 astrocytes **(f)** and N=33-36 neurons **(g)** from 3 independent experiments. **(h)** Corresponding quantification of total GFP (LC3) puncta area per soma area in LLOMe or DMSO treated astrocytes. Horizontal bars represent the means of the biological replicates ± SEM. Shown are p-values from a LME model with Holm’s post hoc correction; N=28-35 astrocytes from 3 independent experiments. **(i)** Corresponding quantification of total LAMP1 puncta copositive for LC3 in LLOMe or DMSO treated astrocytes. Horizontal bars represent the means of the biological replicates ± SEM. Shown are p-values from a LME model with Holm’s post hoc correction; N=26-35 astrocytes from 3 independent experiments. Throughout the figure, neurons are outlined in orange and astrocytes are outlined in blue. Scale bars, 10 µm.

**Supplemental Figure 2.**
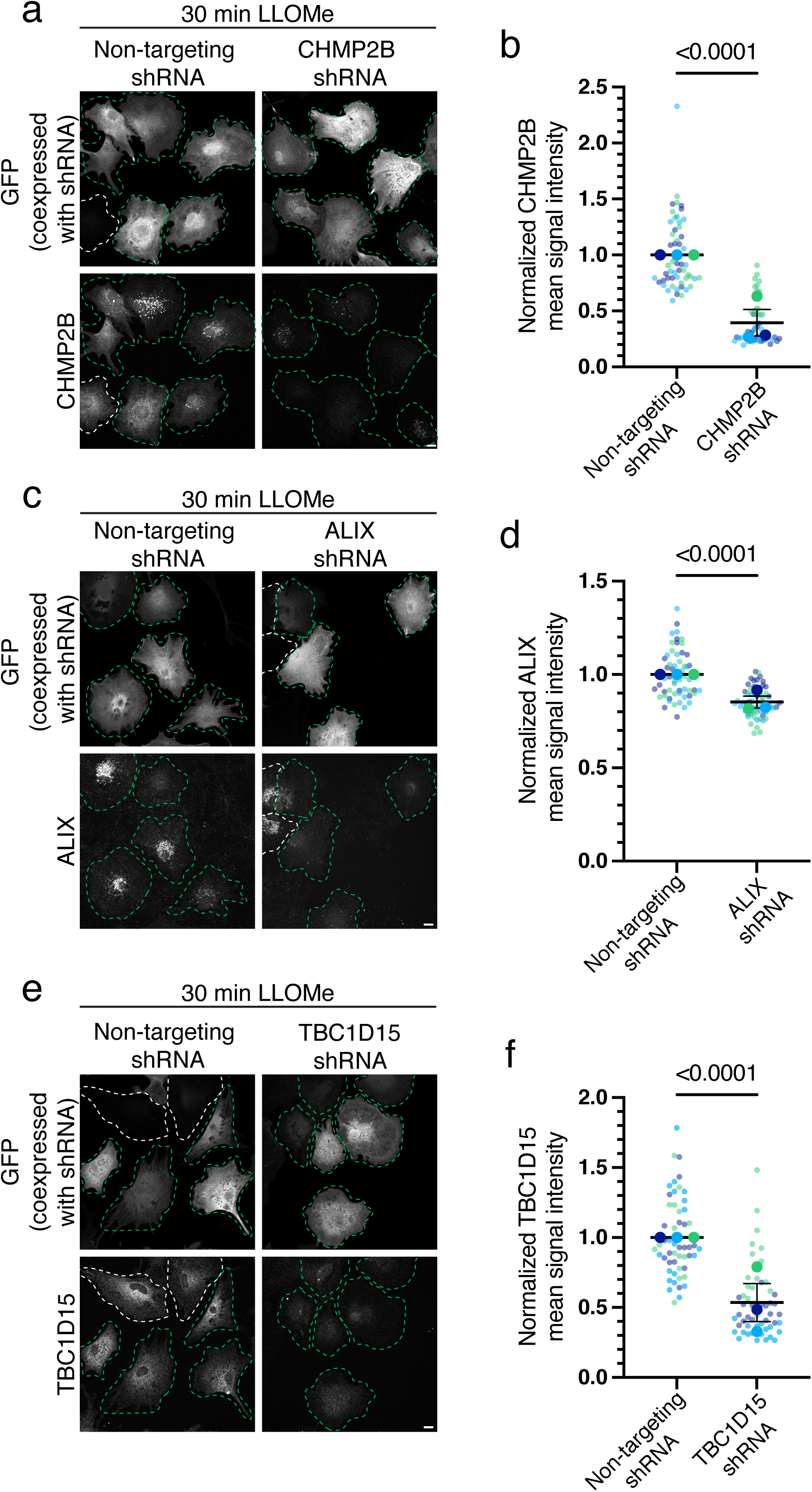
shRNA-mediated knockdown validates antibody specificity for key repair proteins. **(a)** Monocultured cortical astrocytes expressing either a non-targeting shRNA control or a pool of shRNAs to knockdown expression of CHMP2B treated with 1.5 mM LLOMe for 30 minutes and immunostained for GFP (co-expressed with the shRNAs to indicate transduced cells) and CHMP2B. **(b)** Corresponding quantification of normalized CHMP2B mean signal intensity in astrocytes expressing the non-targeting shRNA or the CHMP2B-targeting shRNA. Horizontal bars represent the means of the biological replicates ± SEM. Shown is the p-value from a LME model; N=54-55 astrocytes from 3 independent experiments. **(c)** Monocultured cortical astrocytes expressing either a non-targeting shRNA control or a pool of shRNAs to knockdown expression of ALIX treated with 1.5 mM LLOMe for 30 minutes and immunostained for GFP (co-expressed with the shRNAs to indicate transduced cells) and ALIX. **(d)** Corresponding quantification of normalized ALIX mean signal intensity in astrocytes expressing the non-targeting shRNA or the ALIX-targeting shRNA. Horizontal bars represent the means of the biological replicates ± SEM. Shown is the p-value from a LME model; N=56-58 astrocytes from 3 independent experiments. We note that the ALIX signal intensity in LLOMe-treated control samples is already quite low, which makes it challenging to quantify a large reduction upon knockdown. Nevertheless, ALIX signal is clearly reduced in cells expressing GFP and transduced with the targeting shRNA, and unaffected in cells that are lacking GFP expression and not transduced with the targeting shRNA (**c**). **(e)** Monocultured cortical astrocytes expressing either a non-targeting shRNA or a pool of shRNAs to knockdown expression of TBC1D15 were treated with 1.5 mM LLOMe for 30 minutes and immunostained for GFP (co-expressed with the shRNAs to indicate transduced cells) and TBC1D15. **(f)** Corresponding quantification of normalized TBC1D15 mean signal intensity in astrocytes expressing the non-targeting shRNA or the ALIX-targeting shRNA. Horizontal bars represent the means of the biological replicates ± SEM. Shown is the p-value from a LME model; N=54 astrocytes from 3 independent experiments. Throughout the figure, green outlines denote transduced cells that express the shRNA. White outlines denote non-transduced cells that do not express the shRNA. Scale bars, 10 µm.

**Supplemental Figure 3.**
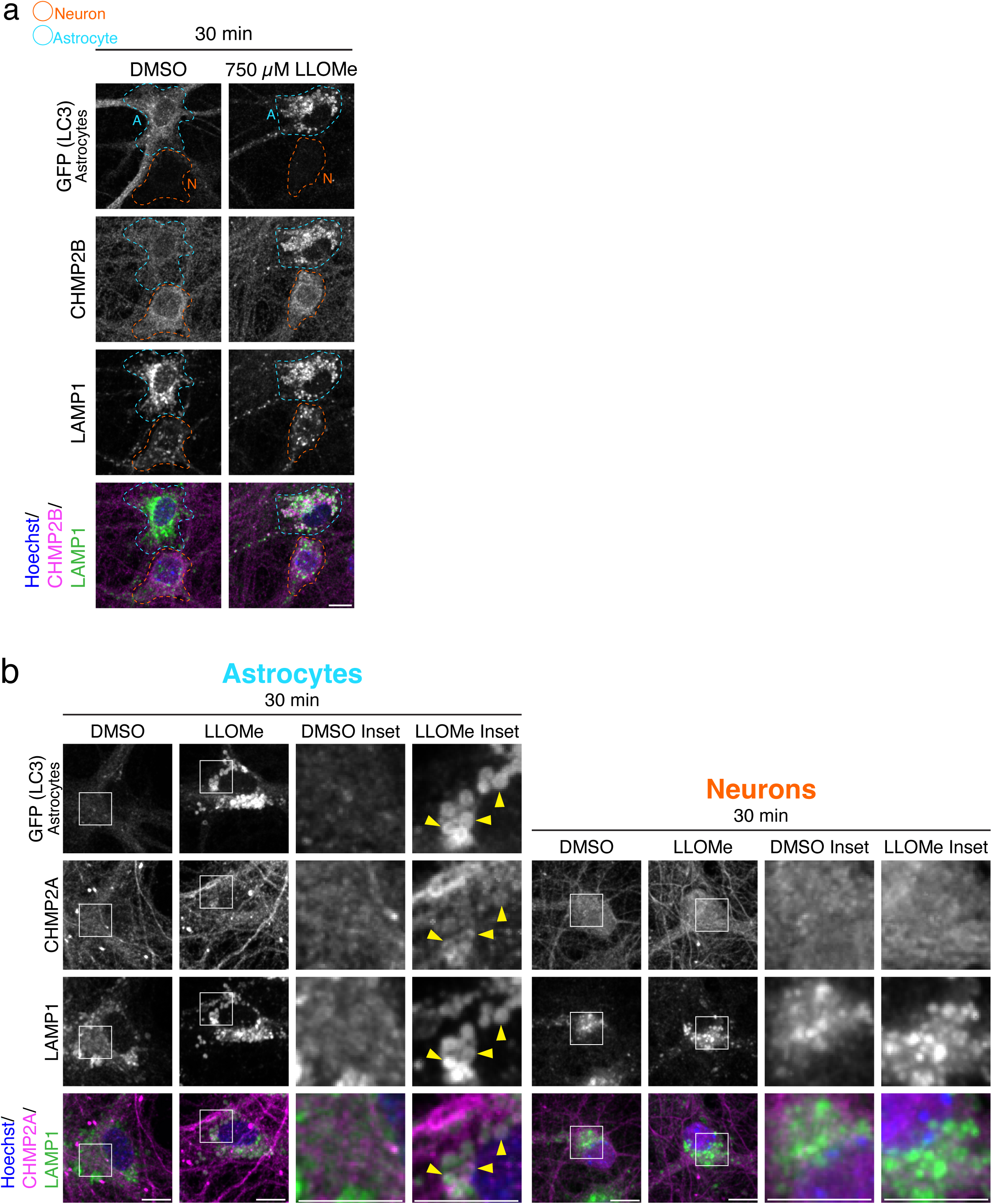
Lower concentration of LLOMe also recruits CHMP2B preferentially in astrocytes compared with neurons. **(a)** Cocultured hippocampal astrocytes and neurons treated with 750 µM LLOMe or equivalent volume DMSO solvent control for 30 minutes and immunostained for GFP (LC3), CHMP2B and LAMP1. Neurons are outlined in orange and astrocytes are outlined in blue. **(b)** Cocultured hippocampal astrocytes and neurons treated with 1.5 mM LLOMe for 30 minutes and immunostained for GFP (LC3), CHMP2A and LAMP1. Insets show zoom-ins of a DMSO and LLOMe treated astrocyte and neuron; location of the inset is indicated with a white box. Yellow arrowheads indicate CHMP2A puncta that overlap with GFP (LC3) puncta and LAMP1 puncta. Scale bars, 10 µm.

**Supplemental Figure 4.**
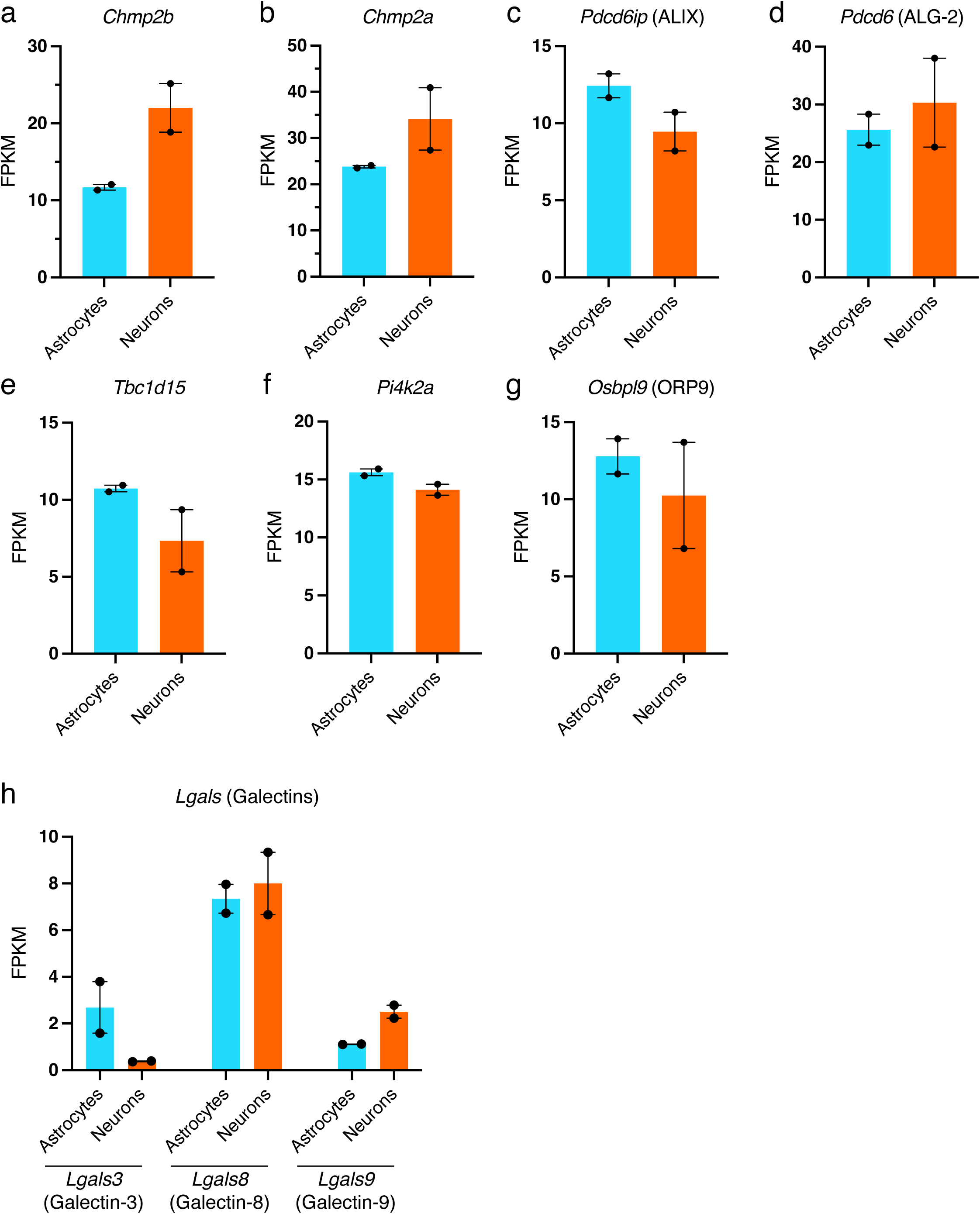
mRNA levels of key lysosomal damage proteins are comparable between astrocytes and neurons. Data extracted from the RNA sequencing database generated by the lab of Dr. Ben Barres showing the mRNA expression profiles of key lysosomal repair proteins in mouse cortical neurons and astrocytes (brainrnaseq.org). **(a-d)** mRNA levels of ESCRT proteins, *Chmp2b* **(a)**, *Chmp2a* **(b)**, *Pdcd6ip* (ALIX) **(c)**, and *Pdcd6* (ALG-2) **(d)** in astrocytes and neurons. **(e)** mRNA levels of *Tbc1d15* in astrocytes and neurons. **(f and g)** mRNA levels of PITT pathway proteins, *Pi4k2a* **(f)** and *Osbpl9* (ORP9) **(g)** in astrocytes and neurons. **(h)** mRNA levels of *Lgals3* (Galectin-3), *Lgals8* (Galectin-8) and *Lgals9* (Galectin-9) in astrocytes and neurons.

**Supplemental Figure 5.**
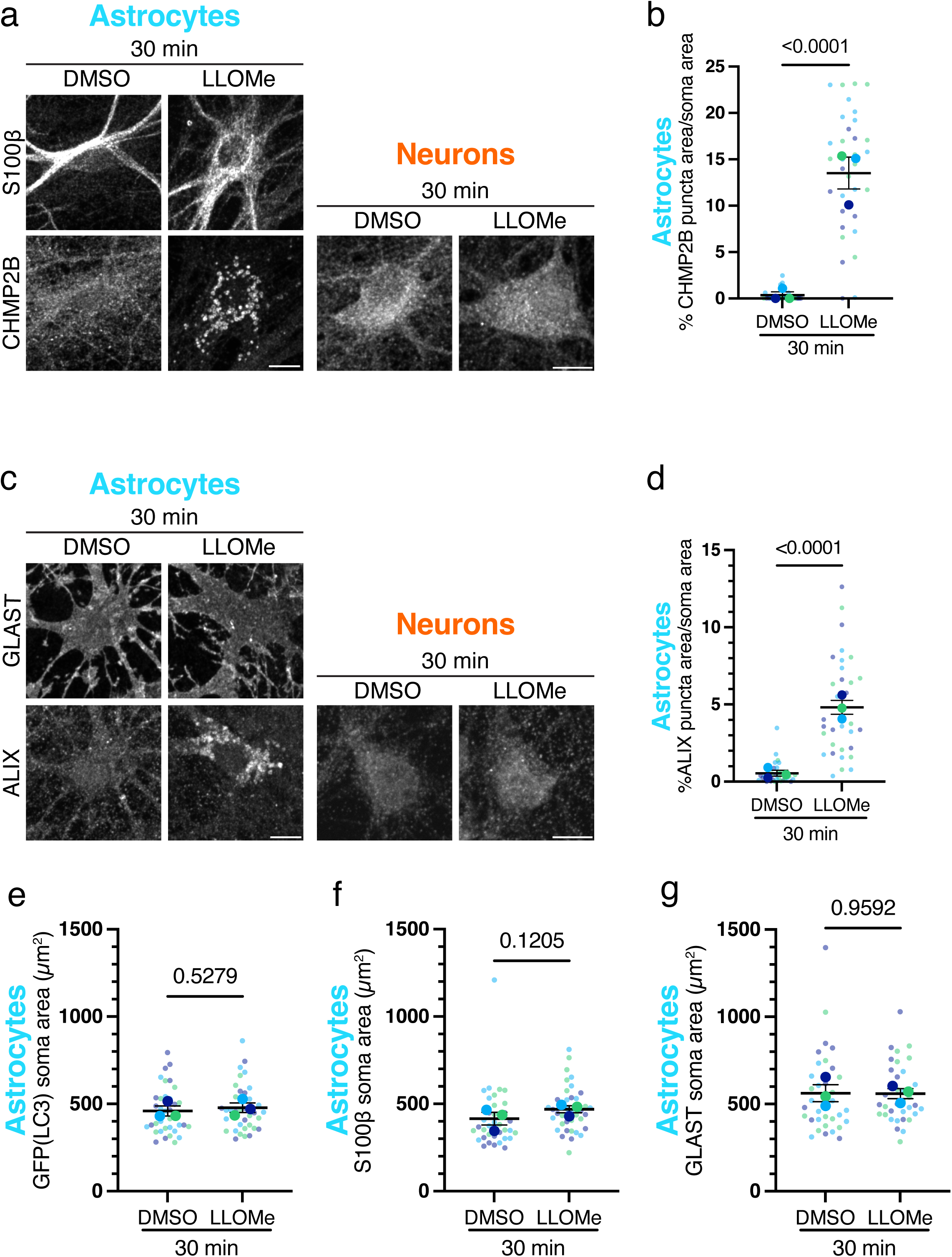
ESCRT recruitment in response to lysosomal damage is conserved in non-transgenic astrocytes. **(a)** Cocultured hippocampal astrocytes and neurons treated with 1.5 mM LLOMe or equivalent volume of DMSO solvent control for 30 minutes and immunostained for S100β and CHMP2B; S100β images in DMSO and LLOMe conditions are not adjusted equally. **(b)** Corresponding quantification of total CHMP2B puncta area normalized to soma area in LLOMe and DMSO treated astrocytes. Horizontal bars represent the means of the biological replicates ± SEM. Shown is the p-value from a LME model; N=33-34 astrocytes from 3 independent experiments. **(c)** Cocultured hippocampal astrocytes and neurons treated with 1.5 mM LLOMe or equivalent volume of DMSO solvent control for 30 minutes and immunostained for GLAST and ALIX. **(d)** Corresponding quantification of total ALIX puncta area normalized to soma area in LLOMe and DMSO treated astrocytes. Horizontal bars represent the means of the biological replicates ± SEM. Shown is the p-value from a LME model; N=31-33 astrocytes from 3 independent experiments. **(e)** Cross-sectional soma area as measured by GFP (LC3) from astrocytes treated with DMSO or LLOMe for 30 minutes. Horizontal bars represent the means of the biological replicates ± SEM. Shown is the p-value from a LME model; N=36-37 astrocytes from 3 independent experiments. **(f)** Cross-sectional soma area as measured by S100β from astrocytes treated with DMSO or LLOMe for 30 minutes. Horizontal bars represent the means of the biological replicates ± SEM. Shown is the p-value from a LME model; N=37-38 astrocytes from 3 independent experiments. **(g)** Cross-sectional soma area as measured by GLAST from astrocytes treated with DMSO or LLOMe for 30 minutes. Horizontal bars represent the means of the biological replicates ± SEM. Shown is the p-value from a LME model; N=36 astrocytes from 3 independent experiments. Scale Bars, 10 µm.

**Supplemental Figure 6.**
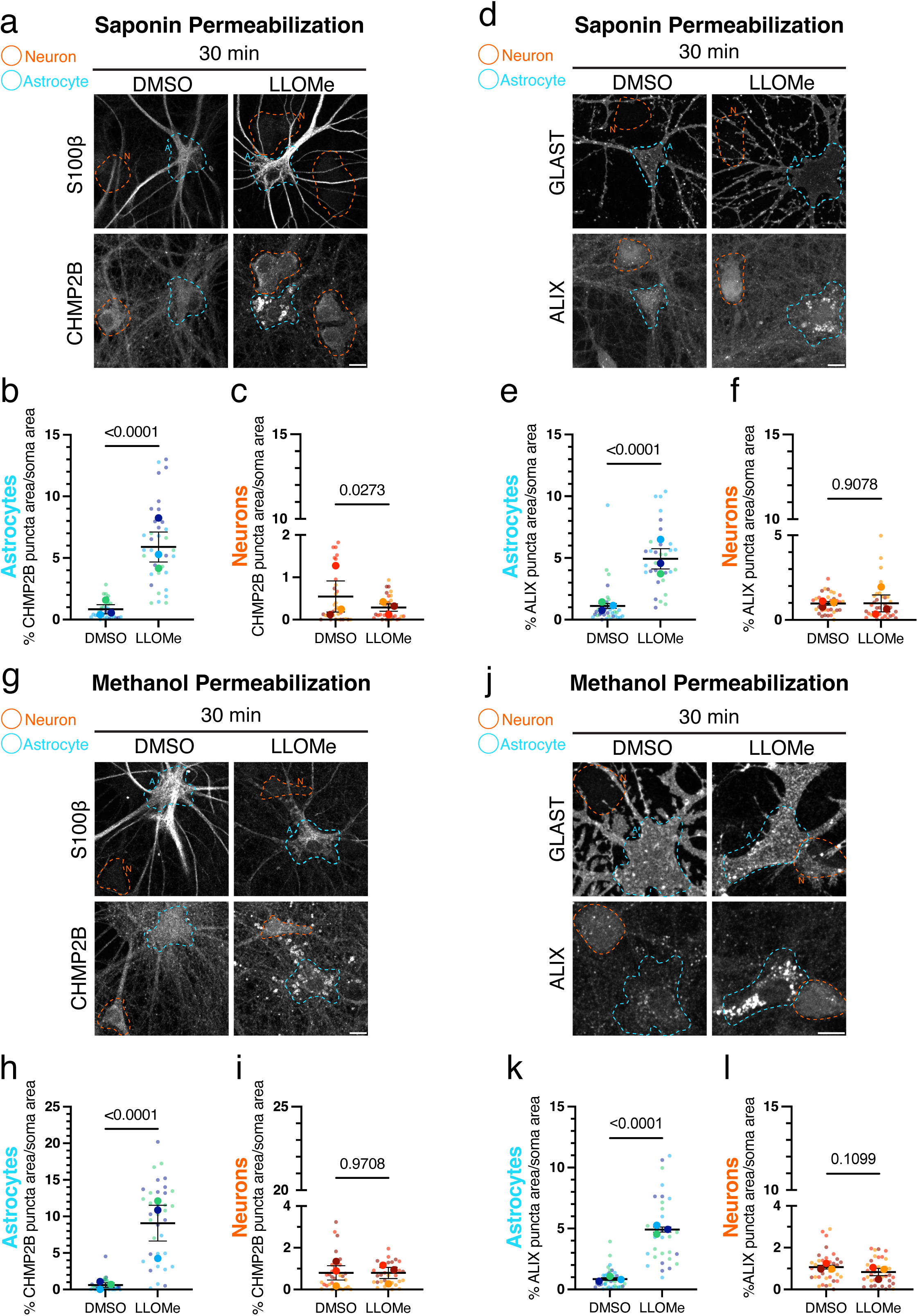
Permeabilization with Saponin or methanol validates preferential ESCRT recruitment in astrocytes observed with Triton X-100 permeabilization in non-transgenic astrocytes. **(a)** Cocultured hippocampal astrocytes and neurons treated with 1.5 mM LLOMe or equivalent volume of DMSO solvent control for 30 minutes, permeabilized with 0.1% Saponin, and immunostained for S100β and CHMP2B. **(b and c)** Corresponding quantification of total CHMP2B puncta area normalized to soma area in LLOMe and DMSO treated astrocytes **(b)** and neurons **(c)**. Horizontal bars represent the means of the biological replicates ± SEM. Shown are p-values from a LME model; N=32-35 astrocytes **(b)** and N=32-34 neurons **(c)** from 3 independent experiments. **(d)** Cocultured hippocampal astrocytes and neurons treated with 1.5 mM LLOMe or equivalent volume of DMSO solvent control for 30 minutes, permeabilized with 0.1% Saponin, and immunostained for GLAST and ALIX. **(e and f)** Corresponding quantification of total ALIX puncta area normalized to soma area in LLOMe and DMSO treated astrocytes **(e)** and neurons **(f)**. Horizontal bars represent the means of the biological replicates ± SEM. Shown are p-values from a LME model; N=33-34 astrocytes **(e)** and N=34-36 neurons **(f)** from 3 independent experiments. **(g)** Cocultured hippocampal astrocytes and neurons treated with 1.5 mM LLOMe or equivalent volume of DMSO solvent control for 30 minutes, permeabilized with 100% methanol, and immunostained for S100β and CHMP2B. S100β images in DMSO and LLOMe conditions are not adjusted equally. **(h and i)** Corresponding quantification of total CHMP2B puncta area normalized to soma area in LLOMe and DMSO treated astrocytes **(h)** and neurons **(i).** Horizontal bars represent the means of the biological replicates ± SEM. Shown are p-values from a LME model; N=35-36 astrocytes **(h)** and N=31-33 neurons **(i)** from 3 independent experiments. **(j)** Cocultured hippocampal astrocytes and neurons treated with 1.5 mM LLOMe or equivalent volume of DMSO solvent control for 30 minutes, permeabilized with 100% methanol, and immunostained for GLAST and ALIX. GLAST images in DMSO and LLOMe conditions are not adjusted equally. **(k and l)** Corresponding quantification of total ALIX puncta area normalized to soma area in LLOMe and DMSO treated astrocytes **(k)** and neurons **(l)**. Horizontal bars represent the means of the biological replicates ± SEM. Shown are p-values from a LME model; N=31-36 astrocytes **(k)** and N=36-37 neurons **(l)** from 3 independent experiments. Throughout the figure, neurons are outlined in orange and astrocytes are outlined in blue. Scale bars, 10 µm.

